# Neuron- and microglia-specific immunoexpression in steroid-independent male sexual behaviour in castrated B6D2F1 male mice

**DOI:** 10.1101/2022.12.08.519640

**Authors:** Vic Shao-Chih Chiang, Sophie Lipkin, Catherine D McCusker, Jin Ho Park

**Author notes:** Corresponding author CHIANG, Vic Shao-Chih, Address Developmental and Brain Sciences, Department of Psychology, University of Massachusetts, Boston, MA, United States.

## Abstract

Sexual behaviour is necessary for procreation for several species and is traditionally viewed to be regulated by sex steroid hormones. However, several species exhibit steroid-independent sexual behaviour, and its molecular understanding is only beginning to be uncovered. The main goal of our experiment was to provide new insight into cell-specific roles that both neuronal and non-neuronal cells may play in steroid-independent male sexual behaviour. Forty B6D2F1 hybrid male mice underwent orchidectomy and were tested for reinstatement of steroid-independent male sexual behaviour after an extended period of social isolation caused by the COVID-19-mandated laboratory shutdown. After 62 weeks post-orchidectomy, 20.59% demonstrated reinstatement of steroid-independent male sexual behaviour (identified as ‘steroid-independent persistent maters’), while 23.53% of the males did not display steroid-independent male sexual behaviour (identified as ‘steroid-independent non-maters’). Using flow cytometry, we compared the preoptic area immunoexpression in NeuN+ neurons and Iba1+ microglia between steroid-independent persistent maters and steroid-independent non-maters (N = 5-6 per group). We found neuronal immunoexpression up-regulated for amyloid precursor protein and androgen receptor, as well as down-regulated for glucocorticoid receptor in steroid-independent persistent maters compared to steroid-independent non-maters. In conjunction, microglial immunoexpression of amyloid precursor protein was up-regulated in steroid-independent persistent maters compared to steroid-independent non-maters. These data suggest there are cell-specific immunoexpression differences, including the role of non-neuronal cells in steroid-independent male sexual behaviour.

## Introduction

Sexual behaviour is integral to many organisms for procreation, social bonding, cooperation, dominance, and even bartering; additionally, it is not limited to the opposite sex and conspecifics (Pelé *et al*., 2017; Monk *et al*., 2019; Addessi & Bourgeois-Gironde, 2020; Steinberg, 2020; Garcia *et al*., 2021; Lemoine *et al*., 2022). The canonical view on how sexual behaviour is regulated involves sex steroid hormones (reviewed in (Wei *et al*., 2021), but several lines of evidence in various species demonstrate the prevalence of steroid-independent male sexual behaviour (SIMSB; reviewed in (Chiang & Park, 2020). One example is the B6D2F1 male mouse, which is a hybrid of C57BL/6J females and DBA/2J males. Approximately one-third of them (~33%) can exhibit gonadal SIMSB for up to four months after orchidectomy (McGill & Tucker, 1964; Manning & Thompson, 1976; Clemens *et al*., 1988). Orchidectomy removes the main source of peripheral steroids, but it’s important to note that other parts of the body, such as the brain, adrenal glands, and sebaceous glands, can still produce steroids (Lolli *et al*., 2017). Therefore, some theories suggest that the sexual behavior observed after castration may be influenced by steroids produced from sources other than the gonads (Sinchak *et al*., 1996). This possibility was ruled out for the adrenal glands, as the removal of the adrenal glands (adrenalectomy) did not eliminate the sexual behavior observed in male mice that had undergone castration (Manning & Thompson, 1976). Even after using drugs that block the androgen receptor, estrogen receptor, and aromatase (an enzyme involved in estrogen production), the sexual behavior in these mice remained unaffected (Park *et al*., 2009).

Previous studies from our lab found molecular differences in the brain between ‘steroid-independent (SI) maters’ (those that demonstrated gonadal SIMSB long after orchidectomy) and ‘SI non-maters’ (Park *et al*., 2010). Through the use of microarray analysis and western blotting, the studies discovered elevated expression of the amyloid precursor protein (App) gene and increased presence of the App protein. Follow-up experiments using transgenic mice that overexpressed APP revealed a higher proportion of mice exhibiting SIMSB compared to wildtype (Park *et al*., 2010). In these experiments, the samples were microdissected from the medial preoptic area (POA) and the bed nucleus of the stria terminalis for downstream assays, as both are integral sites for MSB (reviewed in (Tsuneoka & Funato, 2021). While these generated valuable insights, there were several limitations. One limitation was the heterogeneity of the sample could not be captured, which neglected expression differences in various cell types in the brain and neuron subtypes. With the development of single-cell omics in the past decade, the investigation into the specificity of cell types is much more feasible, substantially propelling our understanding of the neuroscience of social behaviour (Serrano-Saiz & Isogai, 2021; Stein *et al*., 2021). For example, single-cell transcriptomics studies have discovered ~70 different neuronal subtypes in the POA, with specific cell types that are associated with sexual behaviour in males and females (Moffitt *et al*., 2018; Zhang *et al*., 2021). Additionally, an optogenetic study was able to control sexual behaviour by targeting dopamine neurons that projected to the medial POA from the anteroventral and preoptic periventricular area (Zhang *et al*., 2021). However, there remains a lack of knowledge on cell-specific regulations of SIMSB, especially of non-neuronal cells.

In addition to neurons, single-cell sequencing studies uncovered clusters of glial subtypes in brain regions that are crucial for sexual behaviour (Moffitt *et al*., 2018). Of particular interest are microglia, which are resident immune cells in the brain (Lenz *et al*., 2013; Paolicelli *et al*., 2022). Microglia are critical during development to wire neurocircuits required for sexual behaviour (Lenz *et al*., 2013). While it is known that there are other glial types in the brain, such as astrocytes (Blanco-Suárez *et al*., 2017), their specific involvement in sexual behavior has not been established yet. However, further research is needed to explore their potential role in this area before pursuing them as a focus of study. While existing studies do support the involvement of microglia in sexual behaviour, much of this is not understood, especially in terms of adult sexual behaviour. In light of previous findings on SIMSB transcriptomics, it would be of interest to determine to what extent neurons and microglia contributed to the APP up-regulation. Another line of inquiry would be to investigate sex steroid receptors in SIMSB, given how they are paramount for sexual behaviour (reviewed in (Chiang & Park, 2020). Although the preceding microarray analysis did not identify differential gene expression in AR, GR, and ERα at the bulk level (Park *et al*., 2010), further investigation is required to ascertain potential cell-specific variations. Elucidating these cell-specific differences would inform future studies that could test the causal mechanisms of SIMSB.

Here, using the B6D2F1 hybrid male mice, we replicated findings from previous studies that a significant proportion exhibit SIMSB after long-term orchidectomy, but this was at 20.59%, compared to the ~33% in previous studies (Park *et al*., 2009; Templin *et al*., 2019). It is crucial to acknowledge that the current study differs from previous ones because it was conducted during a COVID-19-mandated laboratory shutdown. This shutdown resulted in an extended period of social isolation for all the mice involved in the study. It is well-documented that social isolation can negatively affect the sexual behavior of rodents (Liu *et al*., 2019). For example, in postweaning isolated CD-1 male mice, they exhibited shorter mating duration, longer mount latencies, and longer intromission latencies relative to controls that were not socially isolated (Liu *et al*., 2019). While the effects of extended social isolation on SIMSB have not been directly tested in B6D2F1 hybrid male mice, a study which investigated the role of sexual experience on SIMSB may provide some insight into the potential impact of extended social isolation on SIMSB. Immediately after castration, B6D2F1 hybrid male mice were socially isolated for 12 weeks to withhold sexual experience for these mice (Wee & Clemens, 1989). Interestingly, they found that a significant proportion of the castrated B6D2F1 hybrid male mice were able to demonstrate reinstatement of SIMSB; however, the latencies for mounting, intromission, and ejaculation were longer, as well as the percentage of tests with mounts was lower when compared with males that were given weekly exposure to sexually receptive females (Wee & Clemens, 1989). In our current study, the COVID-19-mandated laboratory shutdown led to our mice being socially isolated for 23 weeks, and all of the mice that had demonstrated SIMSB consistently prior to the shutdown did not demonstrate any MSB on the first sexual behaviour test back. However, after they were provided social exposure, 20.59% of the males demonstrated consistent SIMSB at least 62 weeks post-orchidectomy. Because this current study differed from the previous SIMSB paradigms in our lab, we refer to the males that demonstrated SIMSB in this study as ‘SI persistent maters’ and those that did not as ‘SI non-maters’ to differentiate these mice from prior studies (David, Wyrosdic, & Park, 2022; David, Wyrosdic, Wan, *et al*., 2022). Once SIMSB was reinstated, we then set out to test our original experiment which was to investigate neuron and microglia-specific immunoexpression using flow cytometry in the POA. We hypothesized there would be neuron and microglia-specific expression profiles of APP and steroid receptors that differ between SI persistent maters and SI non-maters.

## Materials & Methods

### Animals

Details of our animal model have been provided previously (Bharadwaj *et al*., 2013). In brief, male B6D2F1 hybrid mice (*Mus musculus;* N = 40, RRID: MGI:5651959) were produced by crossing C57BL/6J females (RRID: IMSR_JAX:000664) with DBA/2J males (RRID: IMSR_JAX:000671). All males were bred and raised in the animal facility at the University of Massachusetts, Boston, weaned at 21 – 28 days, single-sex group-housed until the beginning of experiments (between 50 and 80 d of age), and individually housed afterwards for the rest of the experiments. All mice were housed in the same room, with cages individually ventilated by negative airflow to prevent inter-cage transmission of olfactory cues (Keller *et al*., 2006; McCarthy *et al*., 2017). Animals were given food (Teklad Global Rodent Diet 2016, Harlan Laboratories Inc.) and water *ad libitum* and maintained on a 12:12 light: dark cycle, with lights off at 1200 h EST. All procedures were performed according to the University of Massachusetts, Boston, IACUC (#IACUC2020). All surgeries and sacrifices that were performed were conducted while the animals were anaesthetized with isoflurane, and all efforts were made to minimize suffering including how the mice were handled (Guerra *et al*., 2021). Meloxicam (LKT Labs; 0.5mg/mL in autoclaved distilled water) was given as a post-surgery analgesic at 5mg/kg.

Details of our sexual experience procedure have also been provided previously (McInnis *et al*., 2016). In brief, C57BL/6J females were used as stimulus females for sexual experience and MSB testing. Females were ovariectomized in adulthood after at least one parturition and injected subcutaneously with oestradiol benzoate (MilliporeSigma; 10μg in 0.1 mL sesame oil) 48 h before testing followed by subcutaneous progesterone administration (Millipore Sigma; 400μg in 0.05 mL sesame oil) 3-6h before testing. The females were group-housed in the same colony room as the experimental males. Prior to orchidectomy, B6D2F1 hybrid male subjects were exposed to sexual experience on four distinct occasions by introducing a stimulus female into their home cage and allowing them to remain together overnight.

### Assessment of male sexual behaviour

Details of our male sexual behaviour assessments have been provided previously (Templin *et al*., 2019). In brief, after B6D2F1 hybrid males were surgically castrated with orchidectomy, they were allowed to recover for at least one week before assessing for MSB. The current paradigm differs distinctly from our previous iterations due to COVID-19 where any lab work was suspended for 23 weeks. In this paradigm, the B6D2F1 hybrid males were tested for MSB with stimulus females 11 times over 62 weeks after orchidectomy, and the suspension occurred between tests #2 and #3 (Figure 2). MSB was tested under dim red light at least one hour into the dark phase of a light/dark cycle. All males were habituated at least 30 min before the introduction of the stimulus female in a 17.825.417.8 cm clear Plexiglas testing cage containing the male’s home cage bedding, which had not been changed for at least 48 hours. The test began with the introduction of a hormone-treated stimulus female and videotaped for more than 180 minutes. The videos were scored by an observer for the presence of ejaculatory reflexes. As a result of the COVID-19 situation, the mice were subjected to prolonged social isolation, which could have contributed significantly to the lack of MSB observed in those males who previously exhibited MSB during the second test after castration. Thus, upon returning to the lab at 37 weeks post-castration, each experimental male and stimulus female received social exposure in an attempt to determine if SIMSB could be reinstated after this extended social isolation. Details of the social exposure have been provided previously (Wersinger & Rissman, 2000). In brief, each experimental male and stimulus female received social exposure with a gonad-intact male and female mouse for three consecutive days before MSB. During each exposure, each social stimulus was individually placed in the subject’s home cage for two min. After 11 MSB tests, males were considered as SI persistent maters if they demonstrated ejaculatory reflexes over 40% of all 11 tests, plus over 40% of the last five tests. Males were considered SI non-maters if they never demonstrated any MSB over the 11 tests. To clarify the order of events, a timeline is provided to indicate the sequence of the sexual experience, the COVID-19 mandated laboratory shutdown, the sexual behavior testing, and the subsequent brain extraction step (Figure 1).

**Figure 1.**
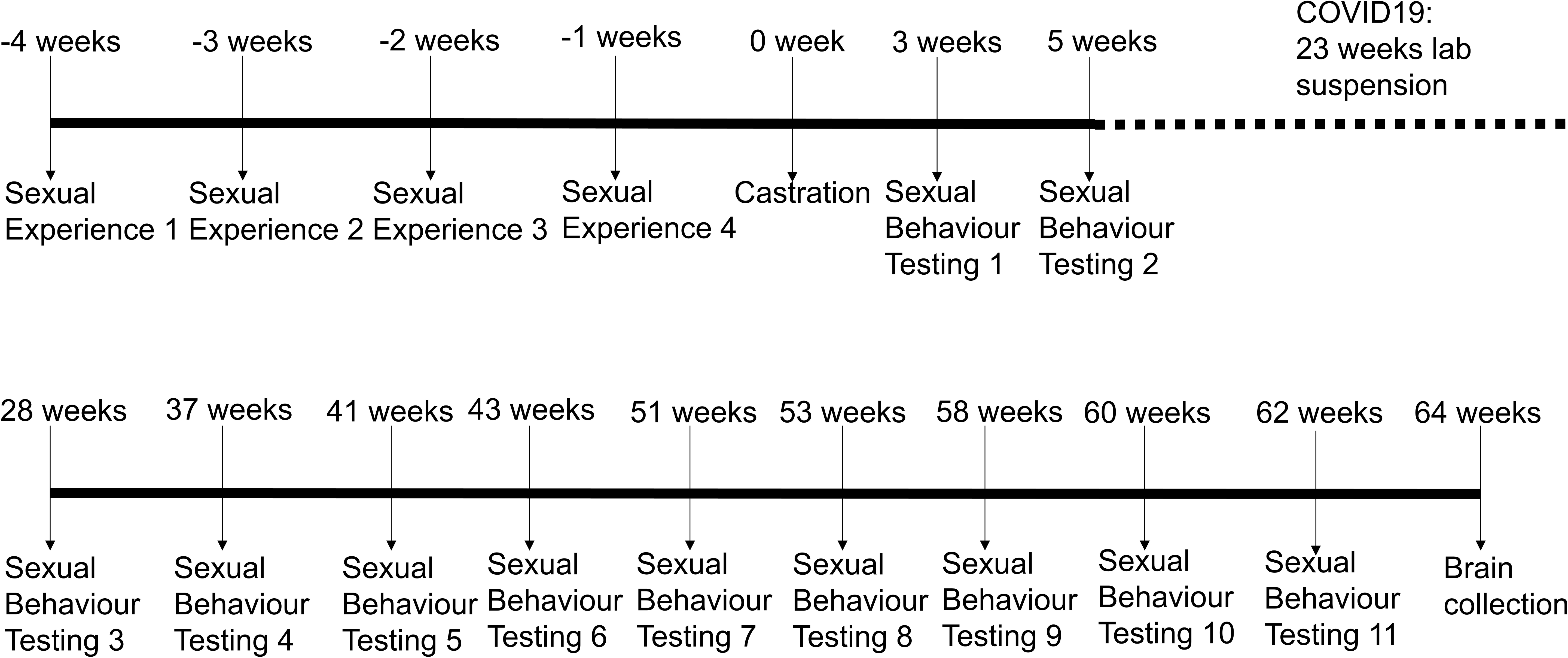
Timeline showing the procedures for the experiment. A chronological representation depicting the sequence of events including sexual experience, castration, sexual behavior testing, laboratory suspension due to the COVID-19 pandemic, and brain collection.

**Figure 2.**
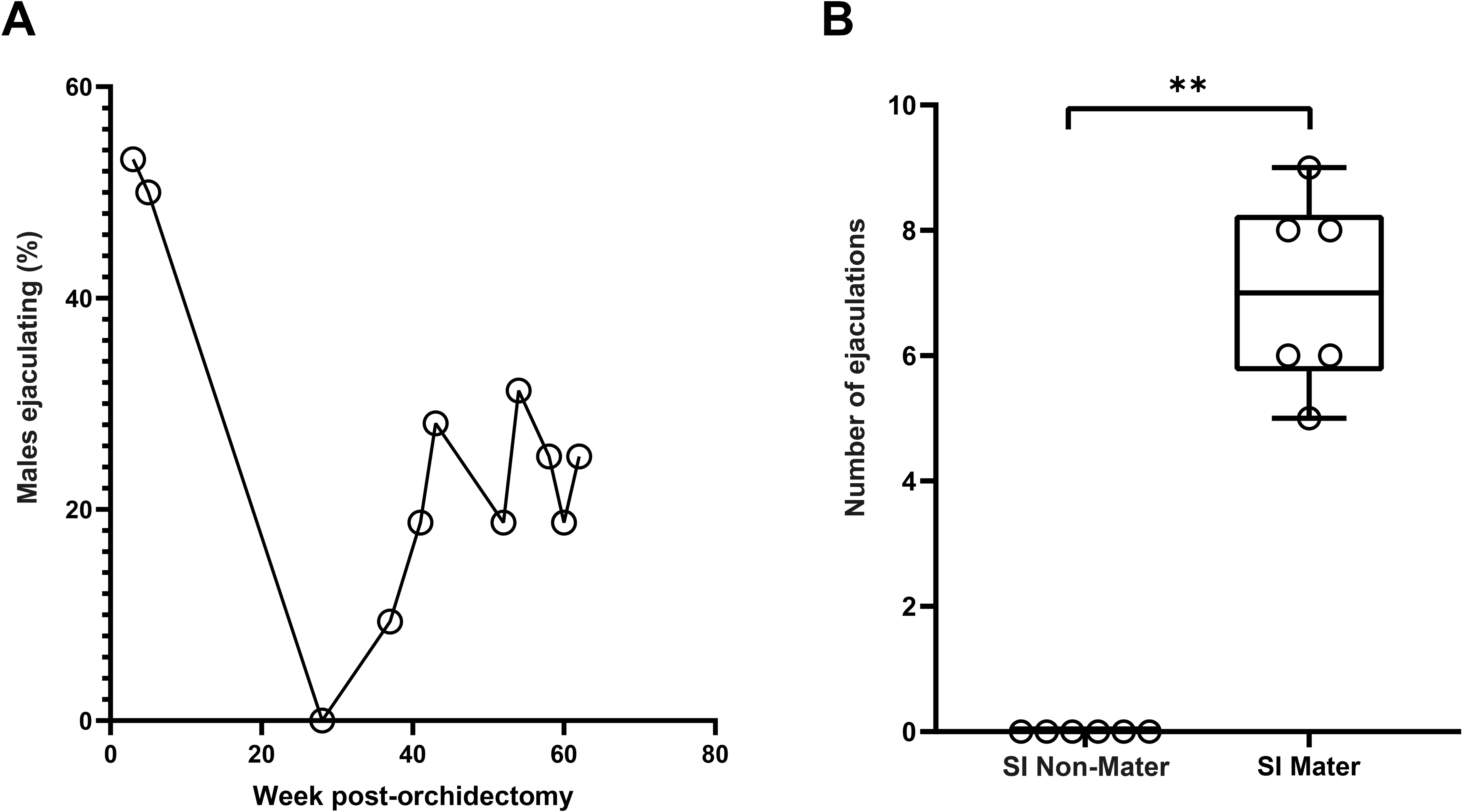
B6D2F1 male mice exhibited male sexual behaviour 62 weeks after orchidectomy. A) Percentage of B6D2F1 male mice exhibited ejaculatory reflex 62 weeks after orchidectomy across 11 tests. B) Quantification of the number of ejaculatory reflexes exhibited by B6D2F1 male mice across 62 weeks after orchidectomy across 11 tests between SI persistent maters and SI non-maters. Data: box and whisker plot showing median with min-max values, data points represent individual samples. Median ejaculatory reflex: SI persistent maters = 7; SI non-maters = 0. Mann-Whitney U nonparametric statistical test, two-tailed, SI persistent mater, N = 6 mice; SI non-mater, N = 6 mice, *p-*value = 0.002, U-value = 0.000, and effect size = 0.892. ** p<0.01; SI, steroid-independent

### Cell isolation

During the last MSB test, one of the six mice that were considered SI-maters did not exhibit ejaculatory reflexes on the last test. This difference within the group would confound our molecular results since we are investigating the sexual phenotype and not ejaculatory reflexes. Therefore, to minimize this potential confounder, we collected brain samples two weeks after the last MSB test (Figure 1). All SI non-maters never exhibited any ejaculatory reflexes post-castration.

Cells were isolated from the adult brain using the Adult Brain Dissociation kit (Miltenyi Biotec, cat # 130-107-677) according to the manufacturer’s instructions, with some modifications made based on recommendations (Jaeger *et al*., 2020) and our own optimization efforts. Given that only four mice could be tractably handled each day, samples were pseudorandomly allocated on different days to include a mix of SI persistent maters and SI non-maters, as well as one extra B6D2F1 hybrid male mouse to provide extra cells for flow cytometry technical controls. Due to the small volume of the target brain region (POA), kidneys were used as carrier cells to improve cell retention during the dissociation. We chose kidneys as it is an easily dissociable organ, and does not substantially contaminate samples with cells expressing markers of nervous system cells, which is common for other easily dissociable organs such as the prostate and the spleen (Pislyagin *et al*., 2017; Haga *et al*., 2019). Mice (N = 16) were rapidly decapitated, and then the brains and kidneys were rapidly extracted and fully immersed into ice-cold Dulbecco’s PBS (Thermo Fisher Scientific, cat# 21600010; free of Mg^2+^ & Ca^2+^; filter-sterilized with 0.22μm filter). The POA was isolated using a brain matrix (Kent Scientific, cat # RBMA-200C) and surgical blades, immersed in ice-cold Dulbecco’s PBS, under a dissection microscope. The POA (preoptic area) sample was supplemented with portions of the kidney to achieve a total sample weight of 400-500 mg. Since any sample that is >400mg provides a sufficient bulk for processing the samples, the amount of kidney added to the POA would not matter. Additionally, the single-cell resolution provided by flow cytometry meant that cells added to the bulk sample would not affect the results, since flow cytometry could gate just the specific brain cells for analysis.

Alongside with kidney, the POA was finely minced in enzyme mix 1. After adding enzyme mix 2, the mixture was incubated in a hybridization chamber (Fisher Scientific, # 13-247-10) with maximum rotation (20min, 35°C). Ensuing that, the mixture was gently triturated with a 20G needle (Becton Dickinson, cat # 305175) and then further incubated (40min, 35°C). Enzymatic digestion was stopped by supplementing to 15mL with DMEM (Gibco, cat# 12800017; added 3.7g/L sodium bicarbonate; adjusted to pH 7.2 – 7.4; filter-sterilized with 0.22μm filter) and added 0.1mg/mL ovomucoid protease inhibitor (Sigma, cat# T2011). The mixture was centrifuged (300 ⍰ *g*, 8min, 4°C) (Thermo Scientific, cat # 50136519) and then filtered through a 70 μm cell strainer (Fisher Scientific, cat # 352350) into a fresh 50mL conical tube, supplementing to 20mL with DMEM. After centrifugation (300 ⍰ *g*, 8 min, 4°C), we carried out the debris removal procedures according to the kit. The suspension was centrifuged with slow acceleration and deceleration rates (900 ⍰ *g*, 15 min, 4°C), and then the supernatant was aspirated. Subsequent to that, the cell pellets were washed by supplementing to 15mL with ice-cold Dulbecco’s PBS and then centrifugation (300 ⍰ *g*, 10 min, 4°C) with full acceleration and brake. Following that, red blood cells were removed according to the kit with the incubation step done using the mini rotator (speed 18, 4°C) (Fisher Scientific, cat # 88861052). To get a sense of the cell viability after these steps, cells were stained with trypan blue (Gibco, cat # 15250061), coverslipped onto a haemocytometer (Hausser Scientific, cat # 02-671-51B), and visualized under a fluorescent cell imager (Bio-Rad, cat # 1450031).

### Staining

Cells were resuspended to a concentration of ~4 – 6 ⍰ 10^6^ cells per ml and stained with 1μL of fixable viability dye, Zombie Aqua (Biolegend; cat# 423101), then incubated in the dark with rotation (room temperature, 30 minutes). The samples were washed with 1mL DMEM, centrifuged (300*g*, 8 min, room temperature) and aspirated until 100 μL remained. Subsequently, the cells were fixed using the Foxp3 / transcription factor staining buffer kit (eBioscience, cat # 00-5521-00) according to the manufacturer’s protocol with several modifications based on the recommendation (Reichard & Asosingh, 2019) and optimization. After fixation, the samples were centrifuged (600 ⍰ *g*, 5 min, room temperature), supernatant aspirated, and then resuspended into 1:1 glycerol (Millipore, cat # 4750-1L): Dulbecco’s PBS (~1 – 2 ⍰ 10^6^ cells per ml). The samples were stored at −20 °C in the dark until permeabilization within two weeks. Before permeabilization, the samples were diluted 10-fold into autoclaved distilled H_2_O and centrifuged (600 ⍰ *g*, 5 min, room temperature) followed by permeabilization according to the manufacturer’s instruction. Afterwards, the samples were divided to incorporate various technical controls, including unstained control, compensation controls, crystallizable fragment receptor blocking control, and fluorescence minus one control. Additionally, two subsamples were prepared from each sample to accommodate two different antibody panels. Each sample and control were then resuspended into 100μL of permeabilization buffer.

Samples and relevant controls were added with 2 μL of crystallizable fragment receptor blocker (Miltenyi Biotec, cat # 130-092-575) and incubated in the dark with rotation (12min, 4°C). The cells were then incubated with the intracellular primary antibody, rabbit anti-Ionized calcium-binding adaptor molecule 1 (Iba1; Fujifilm Wako; cat# 019-19741; RRID: AB_839504; polyclonal; 1:100; 0.5 mg/mL) in the dark with rotation (30min, room temperature). Following that, the cells were washed thrice with 1mL Dulbecco’s DPBS and centrifugation (600 ⍰ *g*, 5 min, room temperature)(Eppendorf, cat # 5424R). Afterwards, the samples and relevant controls were incubated with the secondary antibody, Alexa Fluor 568-conjugated goat anti-rabbit IgG (H+L) (IgG-AF568; Invitrogen; cat# A-11011; RRID:AB_143157; polyclonal; 1:100; 2mg/mL) and the following directly conjugated antibodies: fluorescein isothiocyanate-conjugated rabbit anti-NeuN (NeuN-FITC; Abcam; cat# ab223994; RRID:AB_2892735; monoclonal EPR12763; 1:100; 0.5mg/mL); phycoerythrin-conjugated mouse anti-androgen receptor (AR-PE; R&D Systems; cat# IC5876P; RRID:AB_10890613; monoclonal 523339; 1:10); allophycocyanin-conjugated mouse anti-oestrogen receptor alpha (ERα-APC; Novus Biologicals; cat# NB300-560; RRID:AB_10001032; monoclonal 33; 1:50; 1mg/mL); phycoerythrin-conjugated rabbit anti-amyloid precursor protein (APP-PE; Abcam; cat# ab208744; RRID:AB_2892736; monoclonal Y188; 1:1000; 0.5mg/mL); and allophycocyanin-conjugated mouse anti-glucocorticoid receptor (GR-APC; Invitrogen; cat# 17-6189-82; RRID:AB_2811764; monoclonal BuGR2; 1:200; 0.2mg/mL). These antibodies were selected based on the intended instrument, its spectral signatures, the antigen density to match fluorophore brightness, and available fluorophores for that antibody (Ferrer-Font *et al*., 2020). Supplementary to these, on the basis of prevalent issues of antibodies in terms of specificity, reproducibility and selectivity (Hsu *et al*., 2020), we also selected antibodies from reputable vendors that validated the antibody and cited by existing studies. Anti-Iba1 was validated using mouse retinal whole-mount and cited in an erythro-myeloid progenitors study (Mass *et al*., 2017). Anti-rabbit IgG-AF568 was validated using rat cerebellum and cited in a wind navigation study (Okubo *et al*., 2020). Anti-NeuN-FITC was validated with flow cytometry using PC12 cells and cited in a viral encephalitis study (Michael *et al*., 2020). Anti-AR-PE was validated with flow cytometry using LNCaP cells and cited in a cortical thymic epithelial cell study (Rode & Boehm, 2012). Anti-ERα-APC was validated using human breast tissue and cited in a follicular atresia study (Yao *et al*., 2020). Anti-APP-PE was validated with flow cytometry using SH-SY5Y cells and cited in an adrenergic receptor study (Chena *et al*., 2014). Anti-GR-APC was validated with flow cytometry using mouse splenocytes and cited in an endosomolytic peptide study (Akishiba *et al*., 2017). After staining, samples were subjected to procedures in the Foxp3 / transcription factor staining buffer kit according to the manufacturer’s protocol and then resuspended into Dulbecco’s PBS for flow cytometry analysis.

### Flow cytometry

The fluorescently stained populations (described above) were then analysed using a BD FACSAria Fusion with a 70 μm nozzle. Data were collected using BD FACSDiva software version 6.0 (http://www.bdbiosciences.com/instruments/software/facsdiva/index.jsp; RRID: SCR_001456). A compensation matrix was developed using singly stained and unstained controls of harvested cells. Provisional gates were made using forward and side scatter of light to eliminate debris and non-singlet cells. Dead cells were then excluded using Zombie Aqua as well as Iba1 and NeuN as secondary markers to identify microglial and neuronal populations, respectively. AR, APP, GR and Erα expressions were then analysed. Unstained and fluorescence minus one controls were used for the correct placement of gates. As a benchmark for storage events, more than 30,000 NeuN+ neurons were acquired.

### Flow cytometry bioinformatics

Data were analysed using FlowJo software version 10.6.1 (https://www.flowjo.com/solutions/flowjo; RRID: SCR_008520) following the recommended bioinformatics pipeline (Wang & Brinkman, 2019). The compensated data were quality assessed, and t-transformed using the Logicle function. Subsequently, gating was carried out as described above using contour plots with outliers that show axis labels stating the marker and fluorochrome used, including numbers along the axes. A figure exemplifying the sequential flow cytometry gating strategy can be found in Supplementary Figure 1. Data on AR, APP, GR, and ERα were analysed from the contour plot showing the percentage of neurons or microglia, and from the histogram showing the distribution of the number of neurons or microglia at each fluorescence intensity. The percentage of cells was normalised by dividing the values from a SI persistent mater with a pseudorandomized paired SI non-mater. Pseudorandomization was carried out by listing out all the sample codes for each of the SI persistent maters and SI non-maters in order from the smallest to the largest number. Then, using https://www.random.org/lists/, the sample codes for SI non-maters were pasted to generate the sample codes in random order. This random order was used to pair with the sample codes of SI persistent maters (Supplementary Table 1).

In conjunction with the percentage of cells, we needed a measure of expression intensity. One concern was that flow cytometry fluorescence data were log-normally distributed, which meant that it would have a long-skewed tail that confounds the mean. On the basis of this, we selected the median fluorescent intensity (MdFI) for the measure of expression (O’Neill *et al*., 2013). The MdFI of SI persistent maters were normalized with the pseudorandomized paired SI non-maters (Supplementary Table 1) by the resolution metric. The resolution metric is commonly adopted in flow cytometry experiments (Romano *et al*., 2021; Soh *et al*., 2021; Özgün & Garipcan, 2021; Ditzig *et al*., 2022), because it is able to account for various sources of noise in flow cytometry data (Bhowmick *et al*., 2020). In our experiment, the resolution metric is defined as (MdFI _SI persistent maters_ – MdFI _SI non-maters_)/(rSD _SI persistent maters_ + crSD _SI non-maters_). A higher resolution metric indicates a better separation between the two groups. This takes a similar approach to the ddCT method, which is widely used in qPCR data analysis (Flatschacher *et al*., 2022).

### Preadsorption controls to validate antibodies

Cells were isolated in the same way described in “Cell isolation”, except the samples were from the forebrains of C57BL/6J male mice, without the need for carrier cells. After isolation, cells were stained in the same way described in “Staining” until after the samples were blocked. Each antibody was incubated with fivefold the amount of the respective recombinant protein antigen for one hour at room temperature, and then proceeded with the staining protocol described in “Staining”. Each antibody was single-stained with or without the respective recombinant protein antigen (N = 1 per antibody-antigen combination since this is a validation experiment) (Supplementary Table 2). Flow cytometry was used to collect data as described in “Flow cytometry”, and antibody signals in cells were analysed after dead cells were excluded. Unstained controls were used for the correct placement of gates. As a benchmark for storage events, more than 50,000 cells were acquired. Data were analysed according to “Flow cytometry bioinformatics” except no normalization to the percentage of cells and MdFI were made since the N = 1 for antibody validation did not require statistical analyses.

### Statistical analyses

No statistical methods were used to predetermine sample sizes, but the sample size for flow cytometry experiments are similar to those generally employed in the field (Esk *et al*., 2020; Farmen *et al*., 2021; Sanmarco *et al*., 2021), and sample size for sexual behaviour testing was based on our previous studies (McInnis *et al*., 2016). Biologically independent samples represent six SI persistent maters, and six SI non-maters were used for flow cytometry. Given that there were more than six SI persistent maters and SI non-maters in our mice cohort, we randomly selected six SI persistent maters and six SI non-maters for flow cytometry. Experiments were performed blinded during the annotation of MSB videos to the history of the ejaculatory reflexes of each mouse. Experiments were also blinded to mater status during flow cytometry sample preparation, flow cytometry and bioinformatics analyses. All statistical analyses were performed using SPSS version 28 (https://www.ibm.com/products/spss-statistics; RRID: SCR_019096) and graphs were made with GraphPad Prism version 9.0.0 (http://www.graphpad.com/; RRID: SCR_002798). Formal outlier tests were conducted using Grubb’s test to determine outlying data points, then excluded from the analysis. The data excluded from the analysis is reflected in the changes in the sample size in each result from the original six per group. The final sample size used for each analysis is shown in the figure legends. The outliers from the raw data are provided for reference (Supplementary Table 3), but due to the design of our data analysis, outliers were only excluded after the data were processed as above. Most of the data fulfilled the assumption of Normality based on the Shapiro-Wilks test, without multivariate outliers based on the Mahalanobis distance, but do not fulfil the assumption of the homogeneity of variance based on Levene’s test. Therefore, we proceeded with the non-parametric test, the Mann-Whitney *U* test to compare SI persistent maters and SI non-maters with the threshold for significance set at *p* < 0.05. Statistical details for specific analysis, including exact *N* values and what *N* represents, statistical tests used, central tendency measure, *U* values, *P* values, effect sizes, and all plotted data points, can be found in the figure legends.

To give a sense of the variability of the raw data for the immunoexpression of all molecules measured, we obtained the standard deviation for each molecule (APP, AR, GR, ERα) in neurons and microglia (total N = 8), and used Student’s *T* statistical test to compare between SI persistent maters and SI non-maters with the threshold for significance set at p < 0.05. Since the same animal samples were used for different molecular analyses, we also ran correlation analyses of the immunoexpression of each molecule in neurons and microglia, in terms of how well they correlated with the immunoexpression of all other molecules.

## Results

### B6D2F1 male mice exhibit male sexual behaviour 62 weeks after orchidectomy

At five weeks post-orchidectomy, 50% of the B6D2F1 exhibited ejaculatory reflexes (Figure 2A). Following that, due to COVID-19, we had to suspend any lab work for an extended period. Upon returning to the lab at 28 weeks post-orchidectomy, none of the B6D2F1 exhibited ejaculatory reflexes (Figure 2A). We speculated this could be due to extended social isolation with the absence of physical exposure to other mice, minimal human handling, or ageing (Liu *et al*., 2019; Cho *et al*., 2021; Sokolov *et al*., 2021). In alignment with our commitment to animal welfare and ethical considerations aimed at minimizing animal usage (reviewed in (Smith, 2020), we endeavored to salvage the experiment in question. Social isolation is known to adversely affect MSB (Liu *et al*., 2019), therefore we provided the experimental and stimulus mice with multiple social exposures prior to each sexual behaviour test after 0% of the males showed SIMSB. The repeated sexual behaviour testing and social exposure correlated with an increase in the percentage of males exhibiting ejaculatory reflexes back to levels quasi to previous studies with 28.13% at 43 weeks post-orchidectomy and 25% at 62 weeks post-orchidectomy (Figure 2 A). Previous studies unearthed two distinct sexual phenotypes in these post-orchiectomized B6D2F1 males, namely, the SI maters that consistently exhibit MSB and SI non-maters that do not (Park *et al*., 2009; Bharadwaj *et al*., 2013; Templin *et al*., 2019). In these studies, SI maters were generally defined as individuals who exhibited ejaculatory reflex in at least three out of the last four tests, including the most recent test. On the other hand, SI non-maters were defined as individuals who did not show ejaculatory reflex in the last four tests (Park *et al*., 2009; Bharadwaj *et al*., 2013; Templin *et al*., 2019). Given changes to our paradigm due to COVID-19, our current cohort is distinctly different to previous iterations, and could not utilize the same definition from previous studies. To incorporate the SIMSB reinstatement aspect of this study paradigm, we have labelled the mice that exhibited ejaculatory reflex at least 40% across all 11 tests and at least 40% in the last 5 tests as ‘SI persistent maters’ (seven out of 34 mice were SI persistent maters, 20.59%). The definition of SI non-maters was the same as previous studies, which are those that never exhibited ejaculatory reflexes across all 11 tests (eight out of 34 mice were SI non-maters, 23.53%). We ran statistical analysis on the number of ejaculatory reflexes on the SI persistent maters and SI non-maters we chose for downstream molecular analysis. Our statistical analysis showed SI persistent maters exhibited increased SIMSB compared to SI non-maters (Median ejaculatory reflex: SI persistent maters = 7; SI non-maters = 0. Mann-Whitney U nonparametric statistical test, two-tailed, SI persistent mater, N = 6 mice; SI non-mater, N = 6 mice, *p-*value = 0.002, U-value = 0.000, and effect size = 0.892) (Figure 2 B).

### Steroid-independent persistent maters exhibited altered amyloid precursor protein expression

We next investigated how SIMSB may be regulated at the molecular level in the POA using flow cytometry. Our first target was APP since a previous study found increased APP in SI maters compared to SI non-maters (Park *et al*., 2010). Based on the APP MdFI from the flow cytometry histogram of the SI persistent maters, which was normalized to the SI non-maters using the resolution metrics, we found a decrease in APP immunoexpression in POA neurons (Median MdFI resolution metrics: SI persistent maters neuron = −0.0232; SI non-maters neuron = 0.000. Mann-Whitney U nonparametric statistical test, two-tailed, SI persistent maters neuron, N = 5 mice; SI non-maters neuron, N = 5 mice, *p-*value = 0.003, *U*-value = 0.0000, and effect size = 0.9003) (Figure 3A, 3C). There were no changes to the percentage of neurons expressing APP from the flow cytometry contour plot of SI persistent maters, which was normalized to the SI non-maters (Median normalized percentage of cells: SI persistent maters neuron = 0.9113; SI non-maters neuron = 1.000. Mann-Whitney U nonparametric statistical test, two-tailed, SI persistent maters neuron, N = 6 mice; SI non-maters neuron, N = 6 mice, *p-*value = 0.305, *U*-value = 12.000, and effect size = 0.2962.) (Figure 3B, 3D). This indicates that existing neurons expressing APP reduced APP immunoexpression.

**Figure 3.**
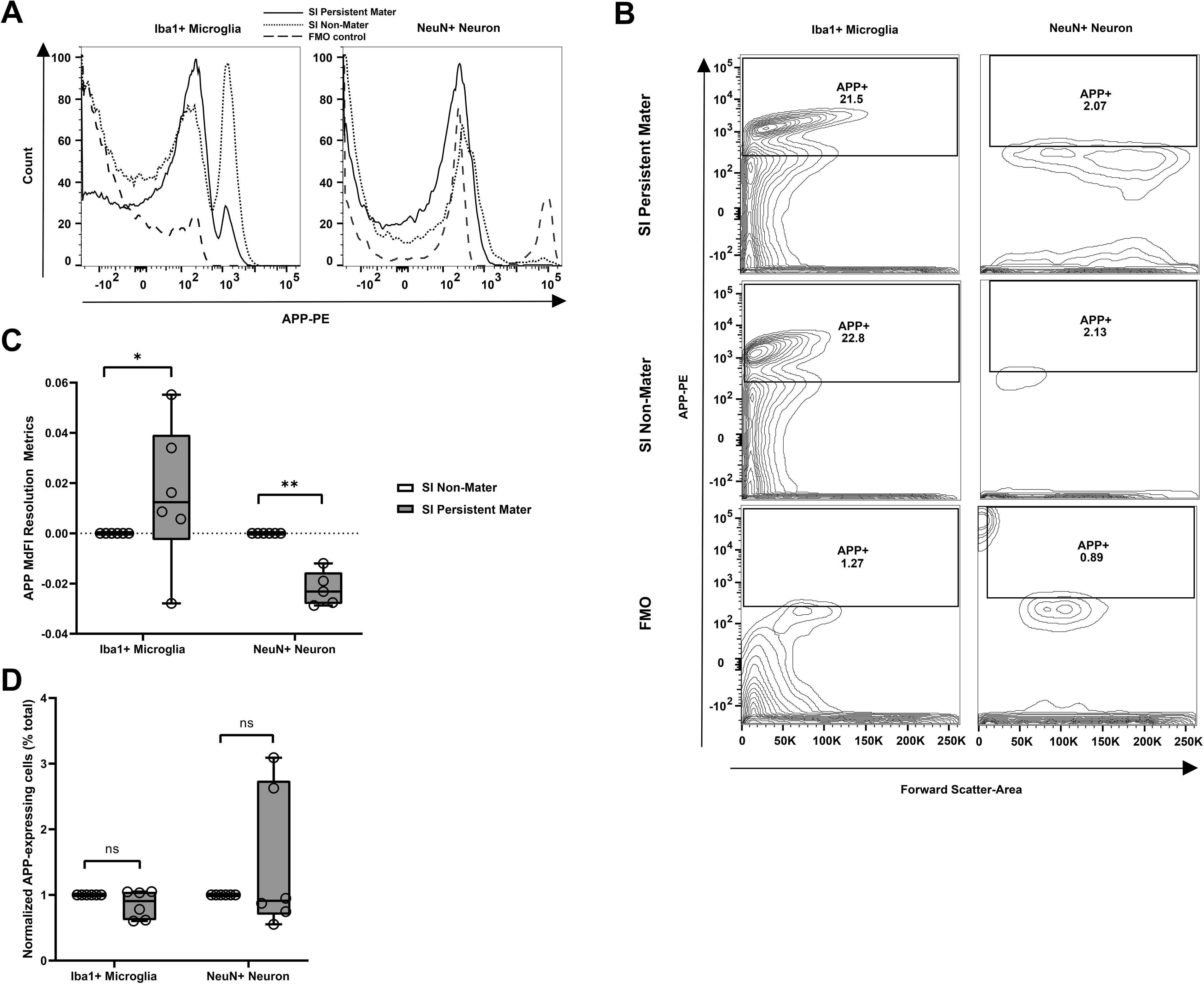
Steroid-independent persistent maters exhibited altered amyloid precursor protein expression in preoptic area microglia and neurons compared to steroid-independent non-maters. A) Representative flow cytometry histogram depicting overlay staining of anti-APP-PE in SI persistent maters (solid line), non-maters (dotted line), and fluorescence minus one control without anti-APP-PE (dashed line), gated on preoptic area microglia (left) or neurons (right). B) Representative flow cytometry contour plot of percentage of APP-expressing microglia or neurons in the preoptic area of SI persistent maters (top), SI non-maters (middle), and fluorescence minus one control without anti-APP-PE (bottom). C) Quantification of the MdFI resolution metrics of APP in the preoptic area neurons and microglia between SI persistent maters and SI non-maters. Data: box and whisker plots showing median with min-max values, data points represent individual samples. Median MdFI resolution metrics: SI persistent maters neurons = −0.0232; SI non-maters neurons = 0.000. Mann-Whitney U nonparametric statistical test, two-tailed, SI persistent maters neuron, N = 5 mice; SI non-maters neurons, N = 5 mice, *p-*value = 0.003, *U*-value = 0.0000, and effect size = 0.9003. Median MdFI resolution metrics: SI persistent maters microglia = 0.0124; SI non-maters microglia = 0.000. Mann-Whitney U nonparametric statistical test, two-tailed, SI persistent maters microglia, N = 6 mice; SI non-maters microglia, N = 6 mice, *p-*value = 0.04, *U*-value = 6.000, and effect size = 0.5921. D) Quantification of the normalized percentage of APP-expressing neurons and microglia in the preoptic area between SI persistent maters and SI non-maters. Data: box and whisker plots showing median with min-max values, data points represent individual samples. Median normalized percentage of cells: SI persistent maters neurons = 0.9113; SI non-maters neurons = 1.000. Mann-Whitney U nonparametric statistical test, two-tailed, SI persistent maters neurons, N = 6 mice; SI non-maters neurons, N = 6 mice, *p-*value = 0.305, *U*-value = 12.000, and effect size = 0.2962. Median normalized percentage of cells: SI persistent maters microglia = 0.9072; SI non-maters microglia = 0.000. Mann-Whitney U nonparametric statistical test, two-tailed, SI persistent maters microglia, N = 6 mice; SI non-maters microglia, N = 6 mice, *p-*value = 1.000, *U*-value = 18.000, and effect size = 0.000. APP, amyloid precursor protein; PE, phycoerythrin; MdFI, median fluorescence intensity; ns, not significant; * p<0.05; ** p<0.01; SI, steroid-independent

There is a paucity of studies investigating non-neuronal cells in the brain in relation to sexual behaviour, therefore, we also investigated APP immunoexpression in Iba1+ microglia. We found an increase in APP expression in Iba1+ microglia of SI persistent maters compared to SI non-maters (Median MdFI resolution metrics: SI persistent maters microglia = 0.0124; SI non-maters microglia = 0.000. Mann-Whitney U nonparametric statistical test, two-tailed, SI persistent maters microglia, N = 6 mice; SI non-maters microglia, N = 6 mice, p-value = 0.04, U-value = 6.000, and effect size = 0.5921) (Figure 3A, 3C). There were no changes in the percentage of APP-expressing microglia in SI persistent maters compared to SI non-maters (Median normalized percentage of cells: SI persistent maters microglia = 0.9072; SI non-maters microglia = 0.000. Mann-Whitney U nonparametric statistical test, two-tailed, SI persistent maters microglia, N = 6 mice; SI non-maters microglia, N = 6 mice, p-value = 1.000, U-value = 18.000, and effect size = 0.000). This indicates that existing microglia expressing APP increased APP immunoexpression.

We ran additional experiments using preadsorption controls as preliminary efforts to help validate the antibodies. All the antibodies were incubated with their respective recombinant protein antigens. For the percentage of cells in preadsorption controls, we found 1.4% NeuN+ cells, and 50.2% with only antibody; 96.4% Iba1+ cells, and 99.5% with only antibody; 98.8% APP+ cells, and 98.9% with only antibody (Supplementary Figure 2; Supplementary Table 4). In terms of the MdFI of the preadsorption controls, these were 68.1 for NeuN, and 236 with only antibody; 88.7 for Iba1, and 232 with only antibody; 107 for APP, and 140 with only antibody (Supplementary Figure 2; Supplementary Table 4). Due to the design of our normalization method, our result graphs did not display the level of variability in the raw data. Therefore, we obtained the standard deviation for all of the molecules measured (AR, APP, GR and ERα), and compared them between the SI persistent maters and SI non-maters, to determine if there were differences in variability. There were no differences in the standard deviation of the percentage of cells between SI persistent maters compared to SI non-maters (Mean standard deviation for the percentage of expressing cells: SI persistent maters = 3.178; SI non-maters = 4.397. Student’s *T* statistical test, two-tailed, SI persistent maters, N = 8 molecules in neurons and microglia; SI non-maters, N = 8 molecules in neurons and microglia, p-value = 0.238)(Supplementary Figures 3). For MdFI, there were no differences in the standard deviation of the MdFI between SI persistent maters compared to SI non-maters (Mean standard deviation for MdFI: SI persistent maters = 67.512; SI non-maters = 68.027. Student’s *T* statistical test, two-tailed, SI persistent maters, N = 8 molecules in neurons and microglia; SI non-maters, N = 8 molecules in neurons and microglia, p-value = 0.990)(Supplementary Figures 3).

### Steroid-independent persistent maters exhibit decreased androgen receptor expression in neurons

The sex steroid receptors, AR and ERα are the cognate receptors of sex steroid hormones, which are paramount for canonical sexual behaviour (reviewed in (Chiang & Park, 2020)). Thus, we compared AR and ERα expression between SI persistent maters and SI non-maters. We discovered a reduced AR expression in NeuN+ neurons of SI persistent maters compared to SI non-maters (Median MdFI resolution metrics: SI persistent maters neuron = −0.0115; SI non-maters neuron = 0.000. Mann-Whitney U nonparametric statistical test, two-tailed, SI persistent maters neuron, N = 6 mice; SI non-maters neuron, N = 6 mice, p-value = 0.04, U-value = 6.0000, and effect size = 0.5921) (Figure 4A, 4C). In addition, we discovered a reduction in the percentage of AR-expressing neuron in SI persistent maters compared to SI non-maters (Median normalized percentage of cells: SI persistent maters neuron = 0.9602; SI non-maters neuron = 1.000. Mann-Whitney U nonparametric statistical test, two-tailed, SI persistent maters neuron, N = 6 mice; SI non-maters neuron, N = 6 mice, p-value = 0.04, U-value = 6.000, and effect size = 0.5921) (Figure 4B, 4D). This indicates that there was an overall reduction of AR expression in SI persistent maters neurons, and this could be related to the reduced neuronal population expressing AR. In terms of non-neuronal cells, Iba1+ microglia did not exhibit differential AR expression in SI persistent maters compared to SI non-maters (Median MdFI resolution metrics: SI persistent maters microglia = −0.007; SI non-maters microglia = 0.000. Mann-Whitney U nonparametric statistical test, two-tailed, SI persistent maters microglia, N = 6 mice; SI non-maters microglia, N = 6 mice, *p-*value = 1.000, *U*-value = 18.000, and effect size = 0.000) (Figure 4A, 4C). In tandem with this, there were no changes in the percentage of Iba1+ neurons expressing AR in SI persistent maters compared to SI non-maters (Median normalized percentage of cells: SI persistent maters microglia = 0.9737; SI non-maters microglia = 0.000. Mann-Whitney U nonparametric statistical test, two-tailed, SI persistent maters microglia, N = 6 mice; SI non-maters microglia, N = 6 mice, *p-*value = 0.305, *U*-value = 12.000, and effect size = 0.2962) (Figure 4B, 4D). By the same token, we examined ERα, but there were no differences detected between SI persistent maters and SI non-maters (Supplementary Figure 2). For the percentage of cells in preadsorption controls, we found 86.6% AR+ cells, and 19.4% with only antibody; 39.9% ERα cells, and 12.9% with only antibody (Supplementary Figure 2; Supplementary Table 4). In terms of the MdFI of the preadsorption controls, these were 68.1 for AR, and 11.6 with only antibody; 585 for ERα, and 293 with only antibody (Supplementary Figure 2; Supplementary Table 4).

**Figure 4.**
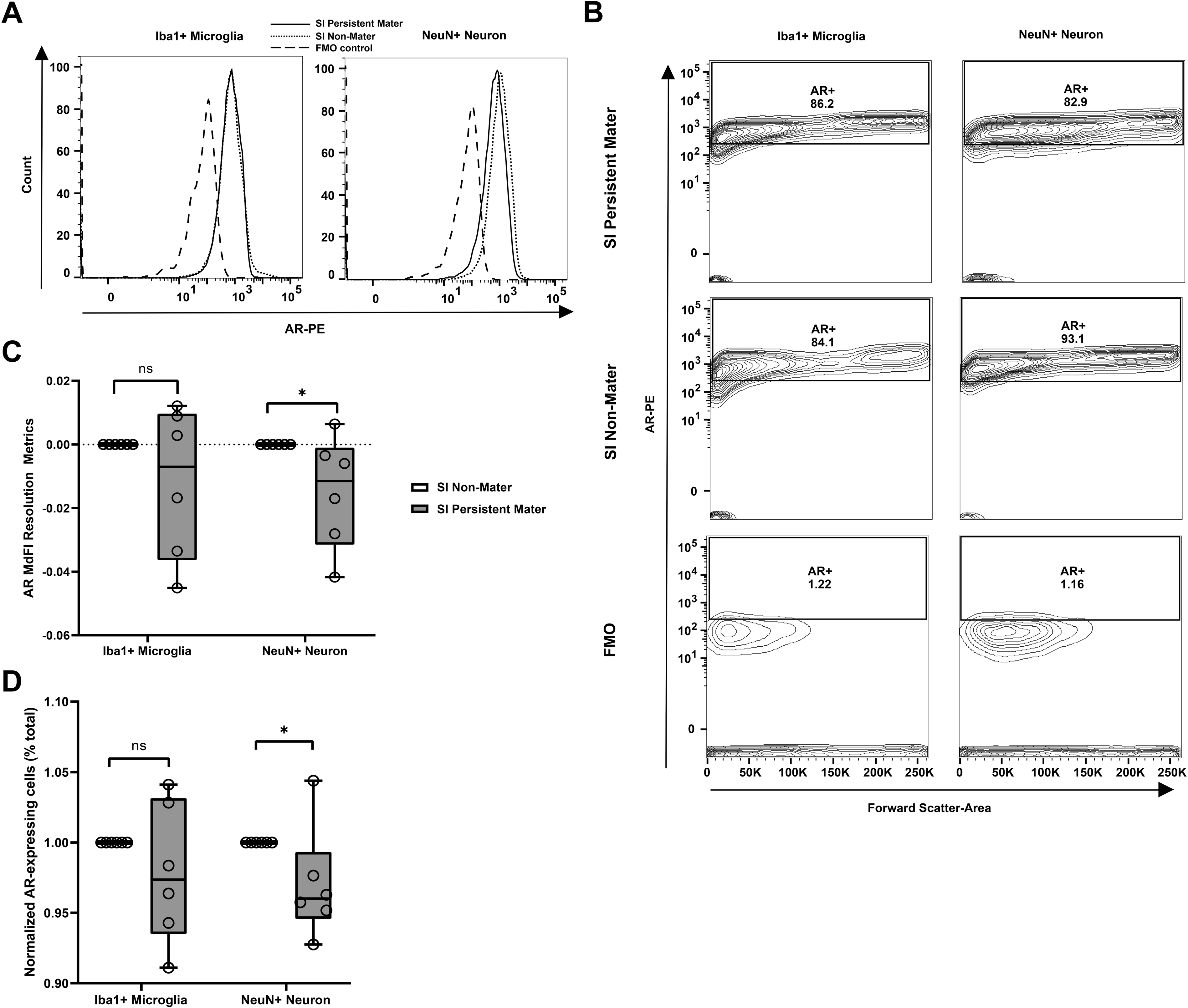
Steroid-independent persistent maters exhibited decreased androgen receptor expression in preoptic area neurons compared to steroid-independent non-maters. A) Representative flow cytometry histogram depicting overlay staining of anti-AR-PE in SI persistent maters (solid line), non-maters (dotted line), and fluorescence minus one control without anti-AR-PE (dashed line), gated on preoptic area microglia (left) or neurons (right). B) Representative flow cytometry contour plot of percentage of AR-expressing microglia or neuron in the preoptic area of SI persistent maters (top), SI non-maters (middle), and fluorescence minus one control without anti-AR-PE (bottom). C) Quantification of the MdFI resolution metrics of AR in the preoptic area neurons and microglia between SI persistent maters and SI non-maters. Data: box and whisker plots showing median with min-max values, data points represent individual samples. Median MdFI resolution metrics: SI persistent maters neurons = −0.0115; SI non-maters neurons = 0.000. Mann-Whitney U nonparametric statistical test, two-tailed, SI persistent maters neurons, N = 6 mice; SI non-maters neurons, N = 6 mice, *p-*value = 0.04, *U*-value = 6.0000, and effect size = 0.5921. Median MdFI resolution metrics: SI-maters microglia = −0.007; SI non-maters microglia = 0.000. Mann-Whitney U nonparametric statistical test, two-tailed, SI-maters microglia, N = 6 mice; SI non-maters microglia, N = 6 mice, *p-*value = 1.000, *U*-value = 18.000, and effect size = 0.000. D) Quantification of the normalized percentage of AR-expressing neurons and microglia in the preoptic area between SI persistent maters and SI non-maters. Data: box and whisker plots showing median with min-max values, data points represent individual samples. Median normalized percentage of cells: SI persistent maters neuron = 0.9602; SI non-maters neuron = 1.000. Mann-Whitney U nonparametric statistical test, two-tailed, SI persistent maters neurons, N = 6 mice; SI non-maters neurons, N = 6 mice, *p-*value = 0.04, *U*-value = 6.000, and effect size = 0.5921. Median normalized percentage of cells: SI persistent maters microglia = 0.9737; SI non-maters microglia = 0.000. Mann-Whitney U nonparametric statistical test, two-tailed, SI persistent maters microglia, N = 6 mice; SI non-maters microglia, N = 6 mice, *p-*value = 0.305, *U*-value = 12.000, and effect size = 0.2962. AR, androgen receptor; PE, phycoerythrin; MdFI, median fluorescence intensity; ns, not significant; * p<0.05; SI, steroid-independent

### Steroid-independent persistent maters exhibit increased glucocorticoid receptor expression in neurons

Lastly, we explored the expression of GR (glucocorticoid receptor) in different cell types, as it shares similarities with ERα and AR (reviewed in (Spaanderman *et al*., 2018). Additionally, glucocorticoids exert their effects through GR, leading to various cellular changes, including the upregulation of glucocorticoid-induced transcript 1 (Kiuchi *et al*., 2019). In a previous study using SIMSB microarray analysis, it was discovered that the expression of glucocorticoid-induced transcript 1 was down-regulated in SI maters compared to SI non-maters (Park *et al*., 2010). Moreover, GR is recognized for its role in regulating sexual behavior under stressful conditions (reviewed in (Ralph *et al*., 2016). We unearthed an increased GR expression in NeuN+ neurons of SI persistent maters compared to SI non-maters (Median MdFI resolution metrics: SI persistent maters neuron = 0.0698; SI non-maters neuron = 0.000. Mann-Whitney U nonparametric statistical test, two-tailed, SI persistent maters neuron, N = 6 mice; SI non-maters neuron, N = 6 mice, p-value = 0.002, U-value = 0.000, and effect size = 0.8883) (Figure 5A, 5C). At the same time, there were no changes in the percentage of GR-expressing neurons in SI persistent maters compared to SI non-maters (Median normalized percentage of cells: SI persistent maters neuron = 0.8824; SI non-maters neuron = 1.000. Mann-Whitney U nonparametric statistical test, two-tailed, SI persistent maters neuron, N = 5 mice; SI non-maters neuron, N = 5 mice, p-value = 0.550, U-value = 12.000, and effect size = 0.18). What this means is the existing neurons acquired higher GR expression. Apropos to comparing SI persistent maters to SI non-maters microglia in the POA, GR expression was unchanged in terms of expression level (Median MdFI resolution metrics: SI persistent maters microglia = 0.0015; SI non-maters microglia = 0.000. Mann-Whitney U nonparametric statistical test, two-tailed, SI persistent maters microglia, N = 6 mice; SI non-maters microglia, N = 6 mice, *p-*value = 0.305, *U*-value = 12.000, and effect size = 0.2962), and percentage of GR-expressing microglia (Median normalized percentage of cells: SI persistent maters microglia = 0.9441; SI non-maters microglia = 0.000. Mann-Whitney U nonparametric statistical test, two-tailed, SI persistent maters microglia, N = 5 mice; SI non-maters microglia, N = 5 mice, *p-*value = 0.073, *U*-value = 6.000, and effect size = 0.54030).

**Figure 5.**
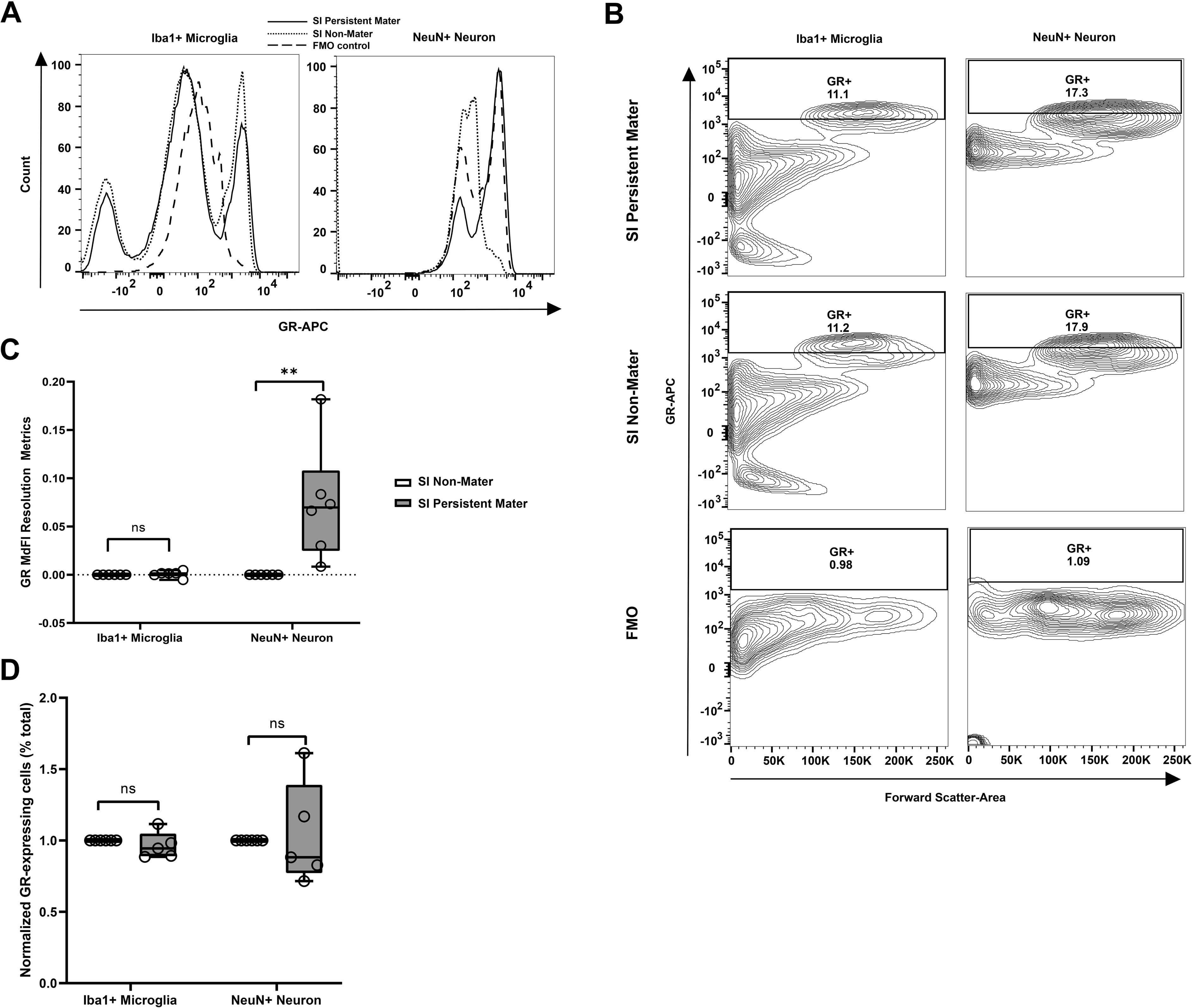
Steroid-independent persistent maters exhibited increased glucocorticoid receptor expression in preoptic area neurons compared to steroid-independent non-maters. A) Representative flow cytometry histogram depicting overlay staining of anti-GR-APC in SI persistent maters (solid line), SI non-maters (dotted line), and fluorescence minus one control without anti-GR-APC (dashed line), gated on preoptic area microglia (left) or neurons (right). B) Representative flow cytometry contour plot of percentage of GR-expressing microglia or neuron in the preoptic area of SI-maters (top), SI non-maters (middle), and fluorescence minus one control without anti-GR-APC (bottom). C) Quantification of the MdFI resolution metrics of GR in the preoptic area neuron and microglia between SI-maters and SI non-maters. Data: box and whisker plots showing median with min-max values, data points represent individual samples. Median MdFI resolution metrics: SI-maters neuron = 0.0698; SI non-maters neuron = 0.000. Mann-Whitney U nonparametric statistical test, two-tailed, SI-maters neuron, N = 6 mice; SI non-maters neuron, N = 6 mice, *p-*value = 0.002, *U*-value = 0.000, and effect size = 0.8883. Median MdFI resolution metrics: SI-maters microglia = 0.0015; SI non-maters microglia = 0.000. Mann-Whitney U nonparametric statistical test, two-tailed, SI-maters microglia, N = 6 mice; SI non-maters microglia, N = 6 mice, *p-*value = 0.305, *U*-value = 12.000, and effect size = 0.2962. D) Quantification of the normalized percentage of GR-expressing neurons and microglia in the preoptic area between SI-maters and SI non-maters. Data: box and whisker plots showing median with min-max values, data points represent individual samples. Median normalized percentage of cells: SI-maters neuron = 0.8824; SI non-maters neuron = 1.000. Mann-Whitney U nonparametric statistical test, two-tailed, SI-maters neuron, N = 5 mice; SI non-maters neuron, N = 5 mice, *p-*value = 0.550, *U*-value = 12.000, and effect size = 0.18. Median normalized percentage of cells: SI-maters microglia = 0.9441; SI non-maters microglia = 0.000. Mann-Whitney U nonparametric statistical test, two-tailed, SI-maters microglia, N = 5 mice; SI non-maters microglia, N = 5 mice, *p-*value = 0.073, *U*-value = 6.000, and effect size = 0.5403. GR, glucocorticoid receptor; APC, allophycocyanin; MdFI, median fluorescence intensity; ns, not significant; ** p<0.01; SI, steroid-independent

For the percentage of cells in preadsorption controls, we found 1.49% GR+ cells, and 1.53% with only the antibody (Supplementary Figure 2; Supplementary Table 4). In terms of the MdFI of the preadsorption controls, these were 585 for GR, and 293 with only the antibody. Aside from the immunoexpression itself, we wanted to see if the immunoexpression of any of the molecules in either the microglia or neurons correlated with other molecules. To do so, we ran correlation analyses for the MdFI, but we did not observe any highly positive (correlation coefficient >0.8) or highly negative (correlation coefficient < −0.8) correlations between different molecules (Supplementary Figure 5).

## Discussion

We show for the first time differential immunoexpression profiles in a cell-specific manner between castrated B6D2F1 hybrid male mice that demonstrate SIMSB relative to those that did not. In terms of neurons, SI persistent maters had down-regulated APP and AR immunoexpression as well as up-regulated GR immunoexpression compared to SI non-maters. For microglia, SI persistent maters demonstrated increased APP immunoexpression compared to SI non-maters. However, there may be caveats with this interpretation due to inconclusive results using the selected antibodies.

Several limitations should be addressed in our study which future studies may help address. The first pertains to the issues of the specificity of antibodies (Hsu *et al*., 2020). For Iba1, GR, and APP, we found no differences in the signals between the antibody-only sample and the preadsorption control. This may indicate these antibodies did not actually bind to the immunogens reported by the vendors. While this may falsify the tested immunogen as the antibody target, our results may still be valid, as it does not eliminate the possibility that other immunogens of the same protein bind to the antibody as well (Holmseth *et al*., 2012; Brownjohn & Ashton, 2014). In the case of ERα and AR, the preadsorption control sample actually demonstrated a higher signal intensity compared to the sample treated with the antibody alone. We suspect for these specific antibodies, there may be heavy complexes formed between the antibody and immunogen, that could not be washed away in our protocol following the commonly cited protocol from Synaptic Systems and R&D Systems. Future studies may need to optimize the protocol for each specific antibody and conduct additional antibody validation experiments to ascertain the validity of results generated from these antibodies.

Another reason for the enhanced antibody signals observed after applying the preadsorption control could be attributed to epitope spreading. Epitope spreading occurs when antigens induce a conformational change in antibodies, causing them to bind to additional epitopes they would not normally bind to (Nguyen *et al*., 2022). This phenomenon has been observed in antitumor immunity and celiac disease (Jin *et al*., 2022; Zhou *et al*., 2022). In our experiment, it is possible that, for certain antibodies, the binding to antigens triggered a conformational change that made the antibodies accessible to other epitopes within our cells. Additionally, antibody aggregation could also contribute to the increased signal observed in some of our preadsorption controls. The binding of the preadsorption antigen may enhance the tendency of antibodies to aggregate (Lai *et al*., 2022). This aligns with the speculation that some heavy complexes may have formed, preventing their removal during washing. Antibody aggregation has been observed in mammalian cell systems and can be induced by free radical oxidation (Zheng *et al*., 2021; Xu *et al*., 2022).

Given the issue with the selected antibodies, we examined relevant literature that used the same clones to see if they encountered similar challenges. For studies that utilized the same NeuN antibody clone (EPR12763) as ours, none of them employed a preadsorption control (Herz *et al*., 2021; Nakamura *et al*., 2021; Xu *et al*., 2021; Georgiou *et al*., 2022). Instead, their focus was on eliminating non-specific signals through a standard blocking step, which we also implemented using the crystallizable fragment blocking control. This pattern was observed for the anti-ERα antibody, using the AER314 clone (Gonçalves *et al*., 2021; Bae *et al*., 2022; Grigoryeva *et al*., 2022), the anti-GR antibody with the BuGR2 clone (Pierce *et al*., 2019; Wofford *et al*., 2020; Præstholm *et al*., 2021; Marchetta *et al*., 2022), and the anti-AR antibody with the clone 523339 (Rode & Boehm, 2012; Pizon *et al*., 2018).

Some of the other antibodies we used presented interesting situations in other studies. Like the previously mentioned antibodies, they lacked a preadsorption control using the same cloned antibodies as ours, and relied on a standard blocking control to address non-specific signals. However, we found two studies that, while lacking a preadsorption control for the same antibodies we used, did include it for one of their other antibodies. This was the case of the Iba-1 antibody (019-19741 from Wako) (Mizuno *et al*., 2006; Ohira *et al*., 2010; Islam *et al*., 2017; Wroblewski *et al*., 2018), where one study included a preadsorption control for anti-progranulin, which successfully eliminated immunoreactivity after preadsorption (Petkau *et al*., 2010). Another antibody with a similar issue was anti-APP, specifically the Y188 clone we used, which only had a preadsorption control for another anti-APP clone they used, known as clone 22C11 (Muresan & Ladescu Muresan, 2015). The authors of these two studies did not provide a specific reason for the selective use of preadsorption controls, but we speculate it could be due to the cost involved in purchasing preadsorption antigens, leading researchers to prioritize which ones to extensively test.

One study did include a preadsorption control for anti-APP antibodies using the Y188 clone we used, which effectively eliminating the signal (Jordà-Siquier *et al*., 2022). They used the APP C-terminal peptide sequence KMQQNGYENPTYKFFEQMQN for the preadsorption control (Jordà-Siquier *et al*., 2022). While our preadsorption control contained this sequence, it was a longer sequence (Suppl Table 2), which might have affected the signal due to the formation of larger immune complexes, as mentioned earlier. Nonetheless, there were still issues with non-specificity, as the same study found antibody signals in white matter tracts in APP gene-deficient mice (Jordà-Siquier *et al*., 2022). Additionally, when co-stained with another APP antibody for the C-terminal clone C1/6.1, the co-localization signal was only 74%, indicating other problems (Jordà-Siquier *et al*., 2022). Furthermore, another study discovered that anti-GR antibodies with the same BuGR2 clone we used produced false signals in thymocytes that were supposed to be GR-deficient (Taves *et al*., 2019).

These studies shed light on several issues regarding antibodies. One concern is the lack of consistent standards for including controls. Another issue is the reliability of antibodies themselves, as demonstrated by a recent study that found inconsistencies with tau antibodies from substantial and concerted efforts with a large research team across several years (Ellis *et al*., 2023). Additionally, due to practical limitations, we were only able to include one sample for each antibody when using preadsorption controls. This prevented us from observing the variability of these experiments, which could have influenced our interpretation of the results.

To enhance the reliability of protein signals, future studies should employ different antibodies to stain for the same protein. It is also crucial to conduct knock-down and overexpression experiments in cell models to validate the antibody signals align with the expected changes. This endeavour requires collaboration between vendors and multiple laboratories, as it may be challenging for individual small labs to carry out alone, as seen in a recent study (Ellis *et al*., 2023). Although these limitations affect the overall conclusions of our study, we believe it serves as valuable preliminary work to explore the involvement of non-neuronal cells in regulating sexual behavior.

The experimental design employed in this study deviates from previous investigations on SIMSB in B6D2F1 hybrid male mice due to an extended laboratory closure resulting from the COVID-19 pandemic. Consequently, the mice were subjected to solitary housing and social isolation for a duration of 23 weeks. However, SIMSB was reinstated and maintained in 20.59% of the mice after repeated social exposure. Our mice model illustrated MSB post-orchidectomy, but the 20.59% of SI persistent maters is less than the one-third observed in previous studies (reviewed in (Chiang & Park, 2020)). The social isolation in our study may not explain this difference fully because a previous study with 12 weeks of social isolation immediately after castration still resulted in 38% of the hybrid males demonstrating SIMSB (Wee & Clemens, 1989). Several differences exist between our study and this previous study. Namely, we had an extended 23 weeks of social isolation, extended testing for 62 weeks post-orchidectomy, and social exposure introduced. In addition, a noteworthy distinction in our study was that the mice utilized for the MSB test were aged 17-18 months, surpassing the conventional breeding age (Moreira *et al*., 2015). Comparatively, the mice in previous studies were around 4 – 6 months of age on their last MSB test (David, Wyrosdic, & Park, 2022; David, Wyrosdic, Wan, *et al*., 2022). Age-related decline in MSB is a known factor that could potentially confound our findings (Wilson *et al*., 2018; Jesús *et al*., 2022). To better understand the contributions of social isolation, social exposure, and aging to the altered percentage of SI persistent maters, future studies should independently investigate these variables. It would be beneficial for these studies to include age-matched controls with intact gonads (with sham operation), as well as mice at the traditional breeding age range of 4-6 months, also with intact gonads (with sham operation) to evaluate the effects of social isolation.

Steroid hormone receptors are pivotal for the development of neurocircuits essential for MSB (Karigo & Deutsch, 2022). We found down-regulated AR and up-regulated GR immunoexpression in POA neurons of SI persistent maters compared to SI non-maters. These findings are inconsistent with the results of a prior microarray analysis, which reported no significant differences in the expression of *Ar* and *Gr* genes between the compared groups (Park *et al*., 2010). Additionally, another previous study demonstrated that AR blockers were ineffective in reducing SIMSB (Park *et al*., 2009). These differences are not surprising, as our approach was cell-specific, as opposed to the bulk approach in previous studies. However, it is important to consider a potential confounding factor, namely the time elapsed between the last MSB test and the collection of brain samples. The duration between these events could impact the molecular profile that was measured. This is because one out of the six SI persistent maters we chose for molecular analysis, did not display ejaculatory reflexes in the last MSB test. Therefore, in our experiments, brain samples were collected two weeks after the last MSB test, to minimize the confound from the ejaculatory reflexes. This was not a problem for SI non-maters, as they never presented any SIMSB post-castration.

There could be an issue with the timing of brain sample collection following the last MSB test. One of the six SI persistent maters did not exhibit ejaculatory reflexes during the final MSB test. This raises the possibility that if additional MSB tests were conducted within the two-week interval, some mice might no longer meet the current definition of SI persistent maters. This quandary arises because different studies have different criteria for classifying mice as SI maters. Instead of a clear distinction between SI maters and SI non-maters, there seems to be a spectrum of SI ‘intermediate’ maters with varying levels of SIMSB observed among cohorts of B6D2F1 hybrid male mice (Park *et al*., 2009). Future studies should develop robust methods to characterize and classify the different SIMSB phenotypic categories. Instead of only considering the ejaculatory reflexes, one option is to include additional measures of sexual behaviour (Heijkoop *et al*., 2018). SIMSB is well-established to differ from the MSB of gonad-intact control without orchidectomy (McGill & Tucker, 1964), but MSB of gonad-intact mice would similarly benefit from this approach to identify nuances in their sexual phenotypes. Traditionally, obtaining multiple sexual behaviour measures is a daunting task, but this is now much more tractable with machine-learning-based behaviour methods (Blau *et al*., 2022; Sun *et al*., 2022)

Nevertheless, we find our comparison to be meaningful as we examined two distinct sexual phenotypes at opposite ends of the spectrum. One end represents mice that never exhibited SIMSB, while the other end represents mice that consistently displayed SIMSB, albeit not in every MSB test. Although the two-week interval between the last MSB test and brain sample collection might affect their status as SI persistent maters, mice showing slightly reduced SIMSB still offer valuable insights into the relevant molecular profiles associated with SIMSB, albeit potentially with a weaker signal.

During development, complementary to neuronal roles to shape sexual behaviour, non-neuronal cells drive spinogenesis essential for sexual behaviour by increasing the production of prostaglandin E2 (Lenz *et al*., 2013, 2018). For microglia, we found up-regulated APP in SI persistent maters compared to SI non-maters, which is in concordance with the previous study that found APP up-regulation (Park *et al*., 2010). However, for neurons, APP was down-regulated. If APP is indeed up-regulated at the global level, what we observed could be that the microglial APP up-regulation may compensate for any down-regulation of neuronal APP. To obtain a more complete picture, future studies can use a multi-method approach to measure APP. This would include repeating the bulk transcriptomics approach for our reinstatement paradigm. Our current study using flow cytometry allowed cell-specific assessment of protein immunoexpression, but future studies can ascertain these results through fluorescence-activated cell sorting followed by western blot on the sorted neurons and microglia. Furthermore, gene expression can be explored using qPCR on the sorted neurons and microglia to validate the congruence between the transcriptional and translational levels.

Another limitation was the normalization method we used. This normalization method has been used in multiple flow cytometry studies (Bhowmick *et al*., 2020; Romano *et al*., 2021; Soh *et al*., 2021; Ditzig *et al*., 2022), and follows a similar logic to the ddCt method used for qPCR (Pabinger *et al*., 2014). In this way, each pair of the comparison group and the control group would combine into only one data point to reflect only their difference. Some argue that this wipes away the variability, but some think that the variability is still reflected in the one data point when the comparison and control group is combined (Pabinger *et al*., 2014). Despite this, we recognize this could be a limitation because there is no consensus on how flow cytometry analysis should be conducted. There are values in future consortiums to identify a gold standard for the field to follow, especially with the burgeoning of bioinformatic techniques that focus on flow cytometry analysis (Cossarizza *et al*., 2021).

From these findings and supporting studies from the literature, we composed a putative signalling pathway for SIMSB (Figure 6). Our targets APP, AR, and GR interact with molecules related to sexual behaviour, namely BDNF (MacPherson, 2017; Brague *et al*., 2018), ERK (Jean *et al*., 2017; Kirouac *et al*., 2017), CREB (Barrot *et al*., 2005; Sun *et al*., 2021), nitric oxide (Melis, 1998; Will *et al*., 2014; Gu *et al*., 2020), glutamate (Numakawa *et al*., 2009), and dopamine 1 receptor (Melis *et al*., 2003; Will *et al*., 2014; Wu *et al*., 2016).They also interact with each other such that AR binds to APP (Takayama *et al*., 2019), AR increases GR transcription (Spaanderman *et al*., 2018) and GR induce secretase processing of APP (Baglietto-Vargas *et al*., 2013). Our correlational analyses did not support this putative signalling pathway, which was expected due to the experimental design being cross-sectional and complicated by the reinstatement of SIMSB. To verify our putative signalling pathway, future *in silico* studies are needed as well as empirical evidence from high-throughput omics that could simultaneously measure millions of molecules at the single-cell resolution. This should include the comparison between the intact control with SIMSB phenotypes, to decipher which molecular changes occurred post-gonadectomy in a mating-dependent manner.

**Figure 6.**
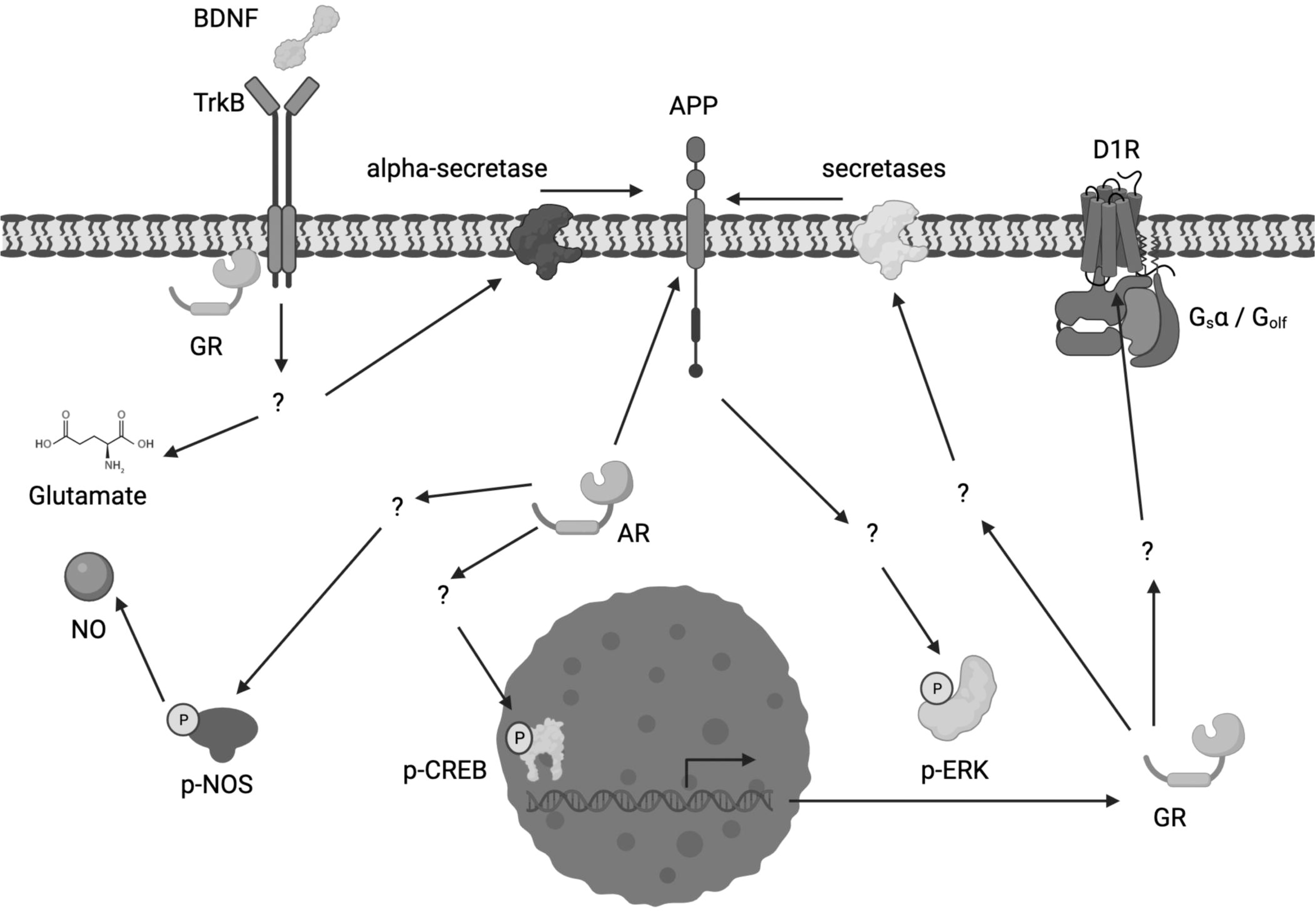
Putative signalling pathway regulating steroid-independent male sexual behaviour. Based on our results, and their molecular targets relevant to sexual behaviour, we constructed a putative signalling pathway that may regulate steroid-independent male sexual behaviour. We observed neuron down-regulation of APP in SI-maters compared to SI non-maters that can occur through BDNF up-regulation of alpha-secretase, which processes APP into smaller fragments (reviewed in (MacPherson, 2017)). Canonical BDNF signalling occurs through binding to TrkB (reviewed in (MacPherson, 2017). APP increase p-ERK (reviewed in (Jean *et al*., 2017). We observed neuron down-regulation of AR in SI-maters compared to SI non-maters, which can directly bind to APP (Takayama *et al*., 2019). AR also increase p-CREB, which drives transcription (Sun *et al*., 2021) and induce p-NOS to produce NO (Gu *et al*., 2020). We observed neuron up-regulation of GR in SI-maters compared to SI non-maters, which can be mediated through AR up-regulation of GR transcription (Spaanderman *et al*., 2018). GR can then induce secretase processing of APP, not limited to alpha-secretase but other secretases including beta and gamma (Baglietto-Vargas *et al*., 2013). Given that secretase processing decreases APP level, the balance of down-regulated AR, and up-regulated GR may result in a reduced APP. GR can be complex with TrkB to trigger glutamate release (Numakawa *et al*., 2009). GR can also up-regulate D1 (Wu *et al*., 2016). BDNF, Brain-Derived Neurotrophic Factor; TrkB, Tropomyosin receptor kinase B; GR, glucocorticoid receptor; APP, amyloid precursor protein; D1R, dopamine 1 receptor; G, Guanine nucleotide-binding proteins; G_s_α, G_s_ alpha subunit; G_olf_, olfactory G; NO, nitric oxide; p-, phosphorylate; NOS, nitric oxide synthase; AR, androgen receptor; ERK, Extracellular signal-regulated kinase;

In summary, we report for the first time cell-specific differential immunoexpression of AR, GR and APP in contrasting sexual phenotypes, defined according to whether or not they exhibited gonadal steroid-independent male sexual behaviour over the long term after orchidectomy and social isolation. Overall, this paves the way to disentangle the environmental context, the phenotypic profile, and the molecular pathways associated with this phenomenon.

## Acknowledgements

We thank Kaylee Wells of the UMass Boston Flow Cytometry Core for support on the BD FACSAria Fusion. We thank Dr Caroline David for assistance with animal husbandry, Benny Wan for assistance with surgeries, and other Park Lab members for general assistance. We thank Renjie Stephen Hu for their help with figures production.

## Author contributions

JHP and VSCC conceived and designed the experiments. VSCC performed the experiments and data analysis with help from SL. CDM supervised the flow cytometry. VSCC created the figures and wrote the manuscript with input from JHP. JHP, CDM and VSCC reviewed and edited the manuscript. JHP supervised the study, provided the resources, and acquired the funding.

## Abbreviations

AF568: Alexa Fluor 568
APC: allophycocyanin
APP: amyloid precursor protein
AR: androgen receptor
ERα: oestrogen receptor alpha
FITC: fluorescein isothiocyanate
GR: glucocorticoid receptor
Iba1: Ionized calcium-binding adaptor molecule 1
MdFI: median fluorescent intensity
MSB: male sexual behaviour
PE: phycoerythrin
POA: preoptic area
SI: steroid-independent

## Data availability

Any data and materials that can be shared will be released via a material transfer agreement. Full access to biological data will be granted to collaborating scientists who are willing to complete a “Data Access Agreement” and a “Data Access Application form” in accordance with General Data Protection Regulation.

## Additional Information

### Supplementary Information

Supplementary material is available at the end of this manuscript.

### Correspondence and requests for materials

Should be addressed to VSC.

### Peer review information

Thanks in advance to the editorial team and reviewers for their contribution to the peer review of this work.

---

**Supplementary Figure 1.**
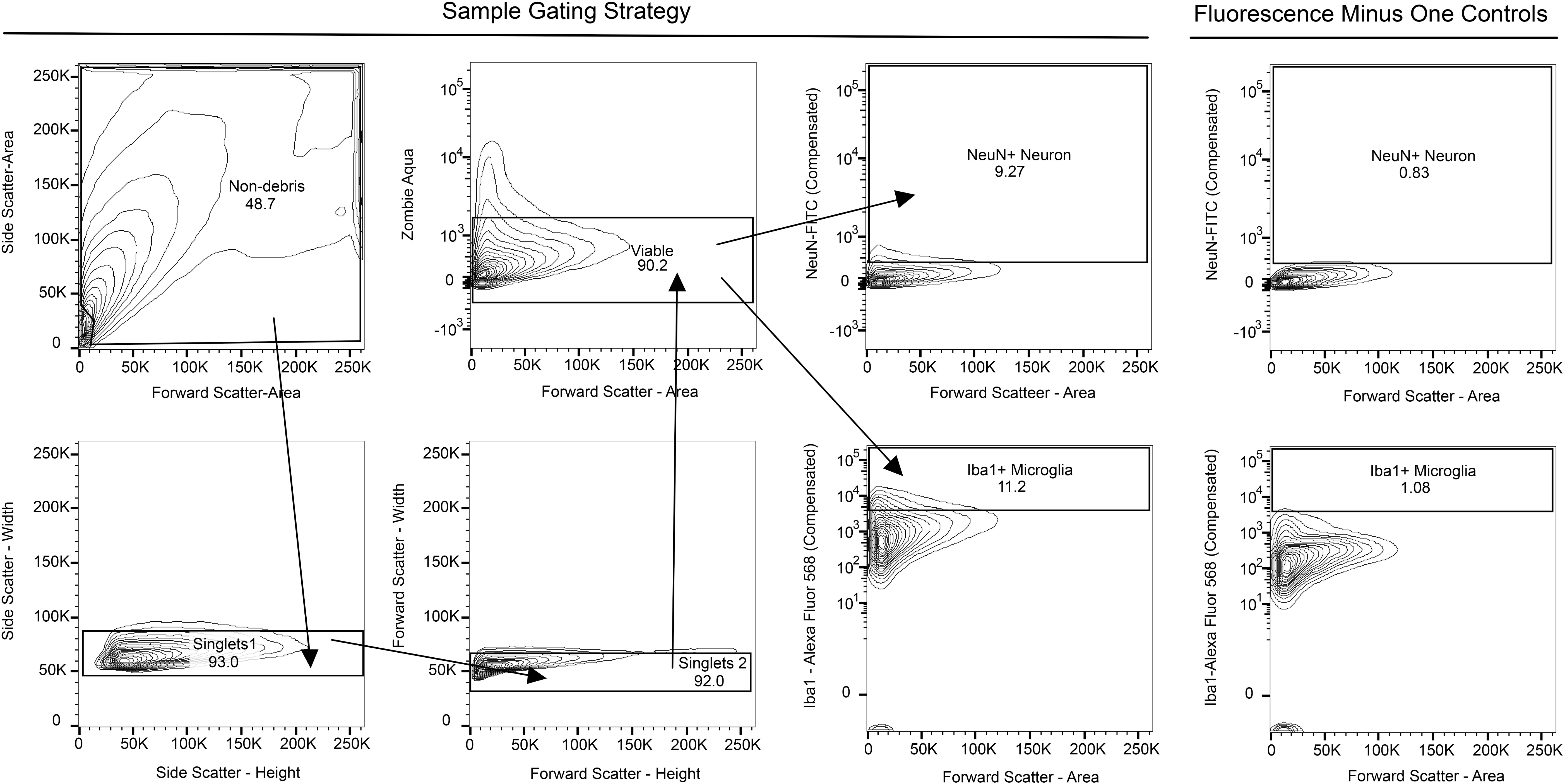
Gating strategy for identifying neuron and microglial populations from flow cytometry data. Representative plots for the gating strategy were used to identify populations of live as well as single microglia and neurons by flow cytometry for analysis. Arrows indicate the pre-gated parent population and the subsequent child population. Forward and side scatter of light were used to eliminate debris, and double exclusion of non-singlet cells by first forward scatter and then side scatter height versus weight. Dead cells were then excluded using fixable Zombie Aqua dye followed by Iba1 and NeuN as secondary markers to identify microglial and neuronal populations, respectively. Candidate molecules expression was then assessed in the Iba1+ and NeuN+ positive population. Unstained and fluorescence minus one controls were used for the correct placement of gates. Iba1, Ionized calcium-binding adaptor molecule 1; NeuN, Hexaribonucleotide Binding Protein-3; FITC, Fluorescein isothiocyanate;

**Supplementary Figure 2.**
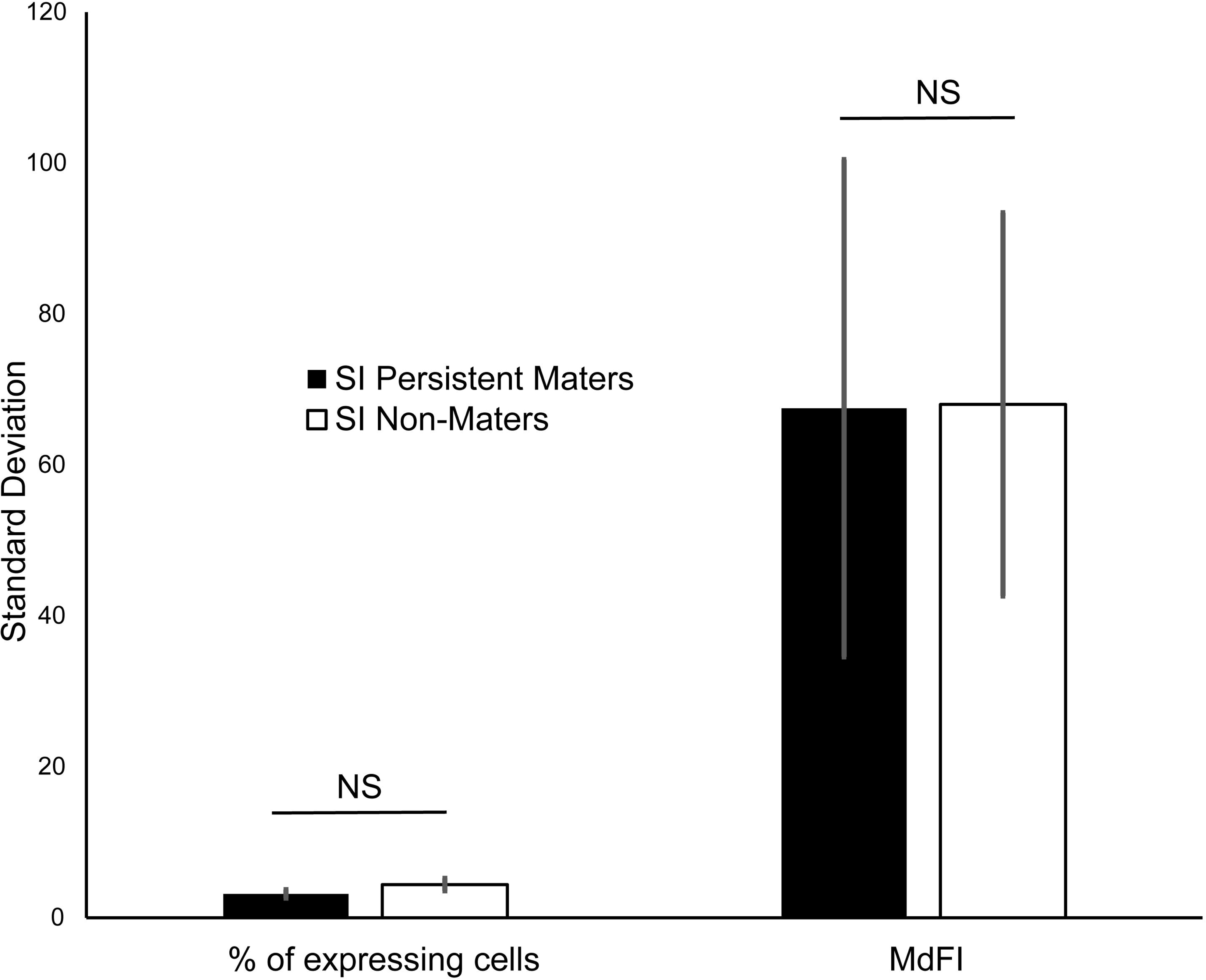
There are no differences in the standard deviations between steroid-independent maters and steroid-independent non-maters. Quantification of the standard deviation for the percentage of cells (left) and MdFI (right) between SI-maters and SI non-maters. Data: bar plots showing the mean percentage of cells (left). Mean standard deviation for the percentage of expressing cells: SI-maters = 3.178; SI non-maters = 4.397. Student’s *T* statistical test, two-tailed, SI-maters, N = 8 molecules in neurons and microglia; SI non-maters, N = 8 molecules in neurons and microglia, p-value = 0.238). Data: bar plots showing mean MdFI (right). Mean standard deviation for MdFI: SI-maters = 67.512; SI non-maters = 68.027. Student’s *T* statistical test, two-tailed, SI-maters, N = 8 molecules in neurons and microglia; SI non-maters, N = 8 molecules in neurons and microglia, p-value = 0.990.

**Supplementary Figure 3.**
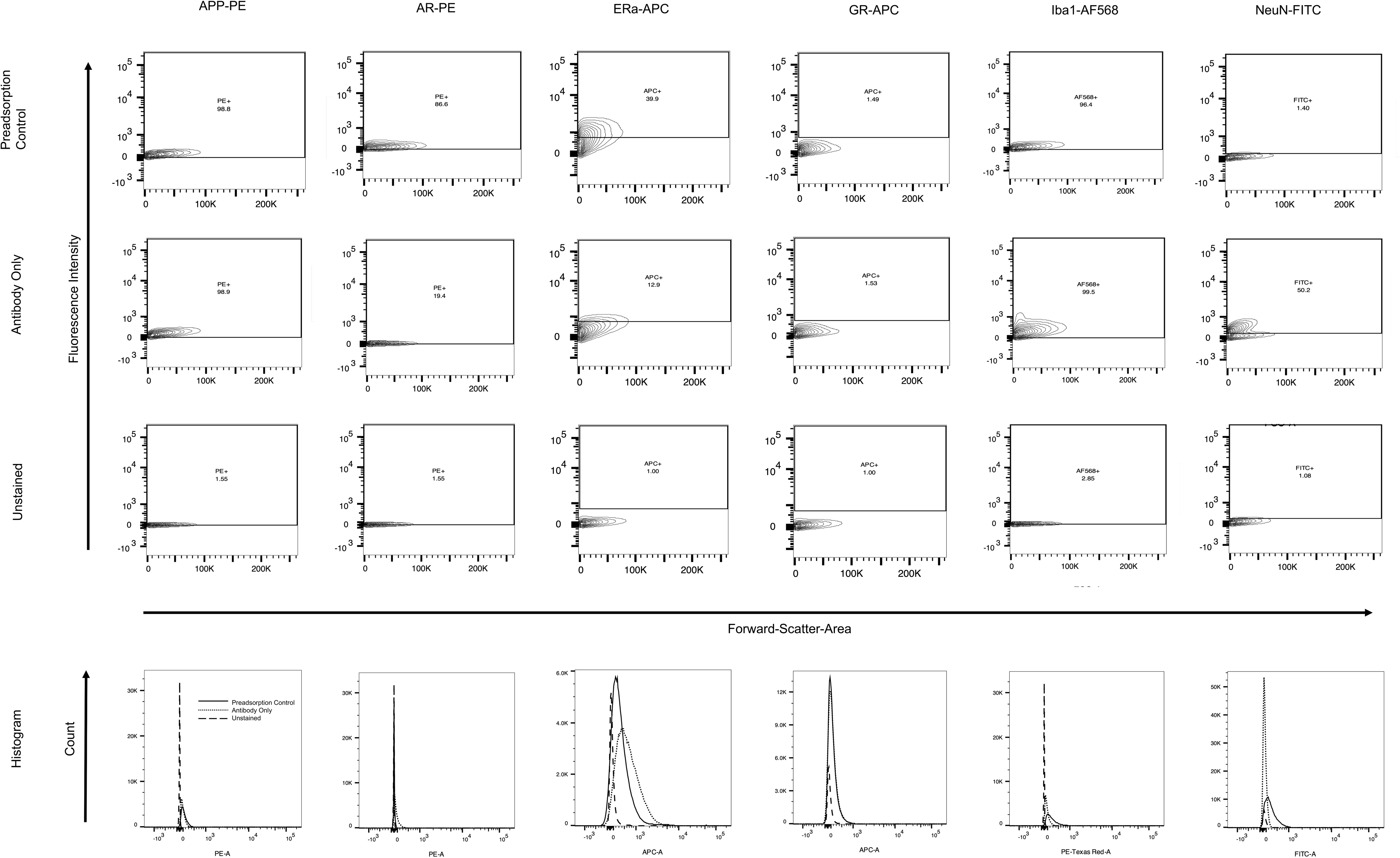
Experiments using preadsorption controls for all the antibodies. Preadsorption controls for all antibodies used: for the forebrain of C57BL/6J male mice, the flow cytometry contour plot (top) of the percentage of cells expressing specific antigens, and histogram (bottom) depicting overlay staining of specific antigens in preadsorption controls (solid line), antibody only (dotted line), and unstained (dashed line). fluorescein isothiocyanate-conjugated rabbit anti-NeuN, NeuN-FITC; phycoerythrin-conjugated mouse anti-androgen receptor, AR-PE; allophycocyanin-conjugated mouse anti-oestrogen receptor alpha, ERα-APC; phycoerythrin-conjugated rabbit anti-amyloid precursor protein, APP-PE; allophycocyanin-conjugated mouse anti-glucocorticoid receptor, GR-APC; anti-Ionized calcium-binding adaptor molecule 1, Iba1; Alexa Fluor 568-conjugated goat anti-rabbit IgG (H+L), AF568;

**Supplementary Figure 4.**
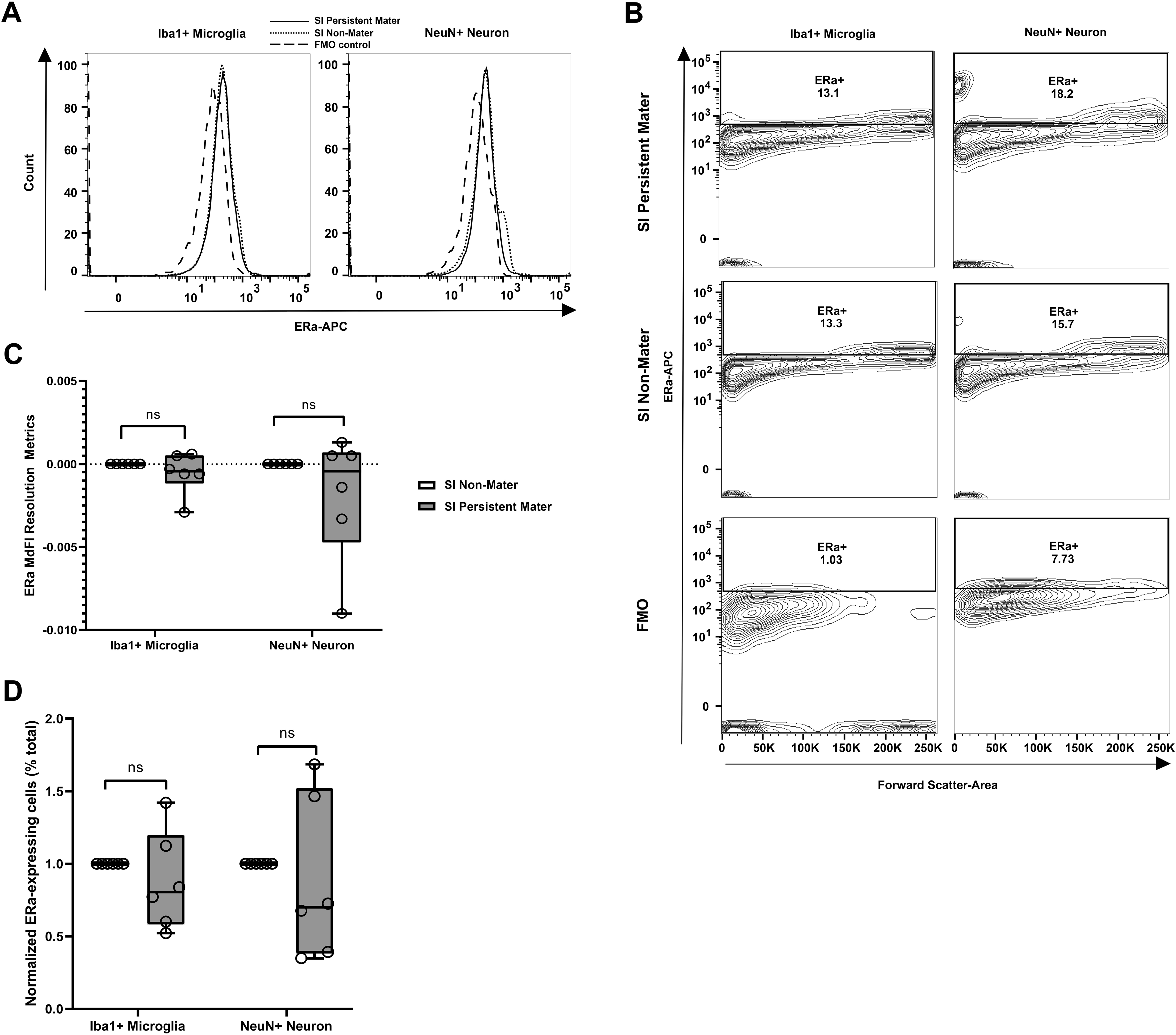
Steroid-independent maters do not differentially express oestrogen receptor alpha in preoptic area neurons compared to steroid-independent non-maters. A) Representative flow cytometry histogram depicting overlay staining of anti-ERα-APC in SI-maters (solid line), non-maters (dotted line), and fluorescence minus one control without anti-ERα-APC (dashed line), gated on preoptic area microglia (left) or neuron (right). B) Representative flow cytometry contour plot of percentage of ERα-expressing microglia or neuron in the preoptic area of SI-maters (top), SI non-maters (middle), and fluorescence minus one control without anti-ERa-APC (bottom). C) Quantification of the MdFI resolution metrics of ERα in the preoptic area neuron and microglia between SI-maters and SI non-maters. Data: box and whisker plots showing median with min-max values, data points represent individual samples. Median MdFI resolution metrics: SI-maters neuron = −0.0005; SI non-maters neuron = 0.000. Mann-Whitney U nonparametric statistical test, two-tailed, SI-maters neuron, N = 6 mice; SI non-maters neuron, N = 6 mice, *p-*value = 1.000, *U*-value = 18.0000, and effect size = 0.000. Median MdFI resolution metrics: SI-maters microglia = −0.0005; SI non-maters microglia = 0.000. Mann-Whitney U nonparametric statistical test, two-tailed, SI-maters microglia, N = 6 mice; SI non-maters microglia, N = 6 mice, *p-*value = 0.305, *U*-value = 12.000, and effect size = 0.2962. D) Quantification of the normalized percentage of ERα-expressing neurons and microglia in the preoptic area between SI maters and SI non-maters. Data: box and whisker plots showing median with min-max values, data points represent individual samples. Median normalized percentage of cells: SI-maters neuron = 0.7014; SI non-maters neuron = 1.000. Mann-Whitney U nonparametric statistical test, two-tailed, SI-maters neuron, N = 6 mice; SI non-maters neuron, N = 6 mice, *p-*value = 0.305, *U*-value = 12.000, and effect size = 0.2962. Median normalized percentage of cells: SI-maters microglia = 0.8053; SI non-maters microglia = 0.000. Mann-Whitney U nonparametric statistical test, two-tailed, SI-maters microglia, N = 6 mice; SI non-maters microglia, N = 6 mice, *p-*value = 0.305, *U*-value = 12.000, and effect size = 0.2962. ERα, oestrogen receptor alpha; APC, allophycocyanin; MdFI, median fluorescence intensity; ns, not significant; SI, steroid-independent

**Supplementary Figure 5.**
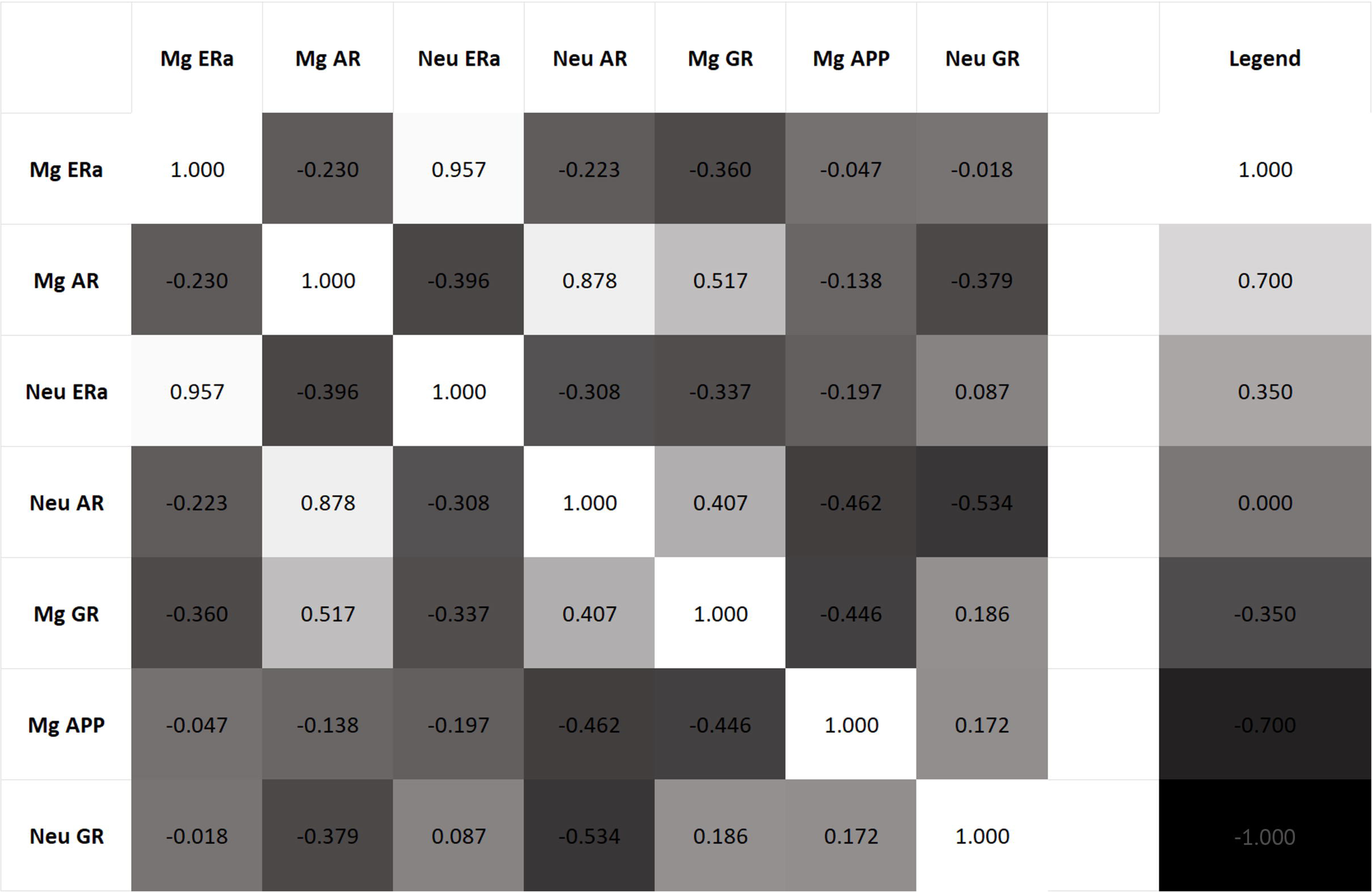
Correlational analyses of androgen receptor, amyloid precursor protein, estrogen receptor alpha, and glucocorticoid receptor in neurons and microglia. Grayscale-coded correlational analyses between MdFI of AR, APP, ERα, and GR in neuron microglia. 1.000 indicate an absolute positive correlation, and −1.000 indicate an absolute negative correlation. AR, androgen receptor; MG, microglia; ERα, estrogen receptor alpha; Neu, neuron; APP, amyloid precursor protein; GR, glucocorticoid receptor.

