## Supplementary material for "Neuron- and microglia-specific immunoexpression in steroid-independent male sexual behaviour in castrated B6D2F1 male mice": Figure Legends & Tables: Tables.docx

**Supplementary Tables**

*Supplementary Table 1. Sample code IDs of steroid-independent persistent maters paired with steroid-independent non-maters.*

| SI Persistent Maters ID | SI Non-Maters Pair |
| --- | --- |
| 2746 | 2770 |
| 2750 | 2777 |
| 2761 | 2755 |
| 2762 | 2768 |
| 2763 | 2774 |
| 2772 | 2760 |

The respective pairs were on the same lines and were generated using random list generators.

SI, steroid-independent.

*Supplementary Table 2. Information on antibodies and the respective recombinant protein antigens used.*

| Antibody | Immunogen | Recombinant Protein Antigen | Sequence of Antigen | Blastp E-value |
| --- | --- | --- | --- | --- |
| Amyloid precursor protein (Abcam, ab208744 | aa 750 to the C-terminus: AVTPEERHLSKMQQNGYENPTYKFFEQMQN | Recombinant Human Amyloid-beta precursor protein(APP),partial (Cusabio, CSB-EP001950HU1) | LKTTQEPLARDPVKLPTTAASTPDAVDKYLETPGDENEHAHFQKAKERLEAKHRERMSQVMREWEEAERQAKNLPKADKKAVIQHFQEKVESLEQEAANERQQLVETHMARVEAMLNDRRRLALENYITALQAVPPRPRHVFNMLKKYVRAEQKDRQHTLKHFEHVRMVDPKKAAQIRSQVMTHLRVIYERMNQSLSLLYNVPAVAEEIQDEVDELLQKEQNYSDDVLANMISEPRISYGNDALMPSLTETKTTVELLPVNGEFSLDDLQPWHSFGADSVPANTENEVEPVDARPAADRGLTTRPGSGLTNIKTEEISEVKMDAEFRHDSGYEVHHQKLVFFAEDVGSNKGAIIGLMVGGVVIATVIVITLVMLKKKQYTSIHHGVVEVDAAVTPEERHLSKMQQNGYENPTYKFFEQMQN | 2.0E-32 |
| Androgen receptor (R&D Systems, IC5876P) | Thr660Gln919: TVSHIEGYECQPIFLNVLEAIEPGVVCAGHDNNQPDSFAALLSSLNELGERQLVHVVKWAKALPGFRNLHVDDQMAVIQYSWMGLMVFAMGWRSFTNVNSRMLYFAPDLVFNEYRMHKSRMYSQCVRMRHLSQEFGWLQITPQEFLCMKALLLFSIIPVDGLKNQKFFDELRMNYIKELDRIIACKRKNPTSCSRRFYQLTKLLDSVQPIARELHQFTFDLLIKSHMVSVDFPEMMAEIISVQVPKILSGKVKPIYFHTQ | Recombinant Human Androgen  receptor(AR),partial (Cusabio, CSB-EP001975HU) | DYYFPPQKTCLICGDEASGCHYGALTCGSCKVFFKRAAEGKQKYLCASRNDCTIDKFRRKNCPSCRLRKCYEAGMTLGARKLKKLGNLKLQEEGEASSTTSPTEETTQKLTVSHIEGYECQPIFLNVLEAIEPGVVCAGHDNNQPDSFAALLSSLNELGERQLVHVVKWAKALPGFRNLHVDDQMAVIQYSWMGLMVFAMGWRSFTNVNSRMLYFAPDLVFNEYRMHKSRMYSQCVRMRHLSQEFGWLQITPQEFLCMKALLLFSIIPVDGLKNQKFFDELRMNYIKELDRIIACKRKNPTSCSRRFYQLTKLLDSVQPIARELHQFTFDLLIKSHMVSVDFPEMMAEIISVQVPKILSGKVKPIYFHT | 0.0E+00 |
| Estrogen receptor alpha (Novus Biologicals, NBP2-48012PCP) | ER purified from cow uterus: No epitope information provided by the company | ER alpha/NR3A1 Recombinant Protein Antigen (Novus Biologicals, NBP1-84826PEP) | LQGEEFVCLKSIILLNSGVYTFLSSTLKSLEEKDHIHRVLDKITDTLIHLMAKAGLTLQQQHQRLAQLLLILSHIRHMSNKGMEHLYSMKCKNVVPLYDLLLEMLDAHRLHAPTSRGGASVEETDQSHLATAGSTSSHSLQKYYIT | N/A |
| Glucocorticoid receptor (Invitrogen, 17-6189-82) | Partially purified rat GR: No epitope information provided by the company | GR/NR3C1 Recombinant Protein Antigen (Novus Biologicals, NBP1-89733PEP) | DSKESLTPGREENPSSVLAQERGDVMDFYKTLRGGATVKVSASSPSLAVASQSDSKQRRLLVDFPKGSVSNAQQPDLSKAVSLSMGLYMGETETKVMGNDLGFPQQGQISLSSGETDLKLLEESIANLNRSTSVPENPKSSASTAVSA | N/A |
| NeuN (Abcam, ab223994) | Within aa1 - 100 (cysteine residue): MAQPYPPAQYPPPPQNGIPAEYAPPPPHPTQDYSGQTPVPTEHGMTLYTPAQTHPEQPGSEASTQPIAGTQTVPQTDEAAQTDSQPLHPSDPTEKQQPKR | RBFOX3/NeuN Recombinant Protein Antigen (Novus Biologicals, NBP1-89821PEP) | PTQDYSGQTPVPTEHGMTLYTPAQTHPEQPGSEASTQPIAGTQTVPQTDEAAQTDSQPLHPSDPTEKQQ | 3.0E-42 |
| Iba1 (Fujifilm, 019-19741) | C-terminus: LGKRSAILKMILMYEEKAREKEKPTGPPAKKAISELP | Recombinant Mouse Allograft inflammatory factor 1(Aif1) (Cusabio, CSB-EP001490HU) | SQSRDLQGGKAFGLLKAQQEERLEGINKQFLDDPKYSNDEDLPSKLEAFKVKYMEFDLNGNGDIDIMSLKRMLEKLGVPKTHLELKRLIREVSSGSEETFSYSDFLRMMLGKRSAILRMILMYEEKNKEHKRPTGPPAKKAISELP | 1.0E-18 |

The information on the antibodies includes the vendor, catalogue number, and the immunogen listed by the vendor. The respective recombinant antigens used with their sequence, vendor and catalogue number. The E-values comparing the antibody immunogen (if available) and the recombinant protein antigen are shown. The BLAST Expect Value (E) describes the number of expected hits of similar quality that could be found just by chance. The smaller the E-number, the better the match.

*Supplementary Table 3. Raw data on the median fluorescence intensity and percentage of viable cells, with Grubb’s outliers test.*

|  | **2746** | **2750** | **2761** | **2762** | **2763** | **2772** | **2760** | **2766** | **2768** | **2770** | **2774** | **2777** |
| --- | --- | --- | --- | --- | --- | --- | --- | --- | --- | --- | --- | --- |
| Phenotype | SI Persistent Mater | SI Persistent Mater | SI Persistent Mater | SI Persistent Mater | SI Persistent Mater | SI Persistent Mater | SI Non-Mater | SI Non-Mater | SI Non-Mater | SI Non-Mater | SI Non-Mater | SI Non-Mater |
| AR+ MG | 82.9 | 87.2 | 87.8 | 90 | 79.3 | 86.2 | 82.8 | 91.1 | 91.5 | 91 | 84.1 | 84.8 |
| ERα+ Neu | 18.2 | 8.68 | 11.4 | 8.73 | 22 | 11.7 | 29.7 | 15.7 | 12.9 | 10.8 | 15 | 24.8 |
| AR+ Neu | 84.6 | 82.9 | 87.9 | 90.9 | 83 | 88.2 | 84.5 | 91.8 | 93.1 | 91.2 | 86.2 | 87.1 |
| ERα+ MG | 18.9 | 13.1 | 11.2 | 11.4 | 18.1 | 13.8 | 26.4 | 14.5 | 13.6 | 13.3 | 16.1 | 21.8 |
| APP+ MG | 18.4 | 17.8 | 20.1 | 18.4 | 21.5 | 17.5 | 29 | 19.2 | 17.5 | 17.8 | 34.8 | 22.8 |
| GR+ MG | 11.1 | 12.4 | 9.63 | 10 | 14.8 | 11.6 | 10.4 | 10.2 | 11.2 | 11.3 | 7.59 | 14 |
| APP+ Neu | 4.02 | 4.08 | 2.06 | 2.07 | 1.66 | 1.86 | 2.13 | 2.17 | 2.77 | 1.53 | 3 | 1.32 |
| GR+ Neu | 16.5 | 25.8 | 12.8 | 17.3 | 24.4 | 18.7 | 16 | 17.9 | 20.9 | 18.7 | 6.12 (Outlier) | 16 |
| Mg ERα | 202 | 194 | 158 | 185 | 173 | 161 | 234 | 179 | 174 | 185 | 184 | 215 |
| Mg AR | 677 | 775 | 734 | 863 | 578 | 715 | 650 | 928 | 848 | 887 | 696 | 715 |
| Neu ERα | 244 | 236 | 200 | 223 | 233 | 207 | 300 | 220 | 217 | 224 | 223 | 279 |
| Neu AR | 728 | 671 | 793 | 953 | 686 | 789 | 744 | 1043 | 975 | 966 | 748 | 818 |
| Mg GR | 50.7 | 59 | 61.4 | 56.6 | 55.7 | 54.4 | 50.1 | 57.4 | 57.2 | 63.1 | 43.3 | 55.6 |
| Mg APP | 11.5 | 43.7 | -16 | 5.09 | -31.5 | -17.2 | -30 | -33.6 | -22.4 | -53.9 | 38.6 (Outlier) | -47 |
| Neu GR | 997 | 1442 | 757 | 909 | 1412 | 1058 | 659 | 721 | 806 | 761 | 683 | 1201 (Outlier) |

The data are shown according to the sample codes of each mouse, with the sexual phenotype information, and the results for each molecule measured in neurons and microglia. The raw data are shown with brackets with “outlier” designated if the value was found to be an outlier in Grubb’s test.

SI, steroid-independent; AR, androgen receptor; MG, microglia; ERα, estrogen receptor alpha; Neu, neuron; APP, amyloid precursor protein; GR, glucocorticoid receptor.

*Supplementary Table 4. Preadsorption control experiment results on the median fluorescence intensity and percentage of viable cells for each antibody.*

| Antibody | Sample | % viable cells | Median fluorescence intensity |
| --- | --- | --- | --- |
| oestrogen receptor alpha | Antibody + antigen | 39.90 | 585.00 |
|  | Antibody only | 12.90 | 293.00 |
|  | Unstained | 1.00 | 61.60 |
| glucocorticoid receptor | Antibody + antigen | 1.49 | 143.00 |
|  | Antibody only | 1.53 | 140.00 |
|  | Unstained | 1.00 | 61.60 |
| NeuN | Antibody + antigen | 1.40 | 68.10 |
|  | Antibody only | 50.20 | 236.00 |
|  | Unstained | 1.08 | 65.50 |
| Ionized calcium-binding adaptor molecule 1 | Antibody + antigen | 96.40 | 88.70 |
|  | Antibody only | 99.50 | 232.00 |
|  | Unstained | 2.85 | 6.42 |
| Amyloid precursor protein | Antibody + antigen | 98.80 | 107.00 |
|  | Antibody only | 98.90 | 140.00 |
|  | Unstained | 1.55 | 5.13 |
| Androgen receptor | Antibody + antigen | 86.60 | 68.10 |
|  | Antibody only | 19.40 | 11.60 |
|  | Unstained | 1.55 | 5.13 |

For each antibody, the result for each antibody + antigen, antibody only and unstained.
